# Temporal persistence of airborne environmental DNA in a natural open-air setting

**DOI:** 10.64898/2026.04.20.719767

**Authors:** Gracie C. Kroos, Kristen Fernandes, Travis Ashcroft, Philip Seddon, Neil J. Gemmell

## Abstract

Airborne environmental DNA (eDNA) is a promising tool for detecting a wide range of terrestrial taxa, including threatened and invasive species, from trace genetic material shed into the air. Yet, application of this technique in species management is constrained by a limited understanding of airborne eDNA ecology, including temporal persistence of signals in open-air environments. We investigated the temporal persistence of airborne eDNA in a natural outdoor setting, using the invasive Bennett’s wallaby *Notamacropus rufogriseus* as a case study. We captured airborne eDNA from a single Bennett’s wallaby carcass, deployed in an area where wallabies are otherwise not present. A total of 180 samples were collected, spanning the period before deploying the carcass, the 11 days it was on site, and for 32 days after its removal, at distances of 1, 10, and 100 metres using both active (fan-assisted) and passive (no fan) collection methods. Although overall detection rates were low, wallaby DNA was detectable up to 100 metres away shortly after the wallaby was introduced to the site and for up to three days after its removal. These findings indicate that airborne eDNA persists only briefly. Actively sampling air using battery-powered fans significantly improved detection rates relative to passive sampling. We demonstrate that airborne eDNA can detect individual organisms in outdoor environments, but reliable detection requires robust sampling and replication to capture rare, transient signals. By revealing how these signals persist over time, our findings provide a framework for optimizing field deployment and for distinguishing remnant DNA from new incursions.

## 1 INTRODUCTION

Airborne environmental DNA (eDNA) is an emerging, noninvasive sampling method capable of detecting species from trace genetic material shed into the air and has strong potential as a tool for terrestrial conservation (Bohmann & Lynggaard, 2023). Air sampling can detect a wide range of taxa, such as birds, mammals, and herpetofauna, as well as invertebrates, plants, and microbial groups such as fungi, including rare, elusive, cryptic, arboreal, or otherwise hard-to-detect species (Johnson et al., 2019; Roger et al., 2022; Garrett et al., 2023a; Lynggaard et al., 2024; Schlegel et al., 2024). While air can capture detailed snapshots of biodiversity (e.g., Sullivan et al. 2025; Tournayre et al. 2025), the pronounced impact of airborne eDNA could be equally realised through deliberate, targeted deployment, for example, at-risk species management (Moloney et al. 2025) or to detect invasive species (e.g., Trujillo-González et al., 2022; Kroos et al., 2026).

Whilst airborne eDNA represents a promising tool for detecting rare taxa such as threatened or invasive species, key knowledge gaps currently hinder its application. Our understanding of eDNA ecology in the air, including signal accumulation, persistence and degradation dynamics, is lacking (Newton et al., 2025; Zhao & Andermann, 2026). In particular, the temporal persistence of airborne eDNA signals in open, natural environments remains poorly understood (Johnson & Barnes, 2024; Tulloch et al., 2025). Yet, this information is critical for optimising field deployment strategies, including sampling duration, and for defining the types of ecological questions that airborne eDNA can reliably address. Without knowledge of signal persistence, it is unclear whether detections reflect recent animal activity or long-term residual signals from past animal presence. This distinction is critical for useful applications such as in invasive species management to differentiate remnant DNA following successful animal removal, from detections associated with new incursions (Dunker et al., 2016).

Degradation of eDNA in aquatic systems is known to occur almost immediately following deposition into the environment through abiotic factors such as light, as well as microbial activity (Barnes & Turner, 2016). Degradation rates increase with higher temperatures (Kasai et al. 2020) and extreme pH (Seymour et al. 2018). In marine and freshwater systems, the signal generated by eDNA often disappears within 4-7 days, and rarely persists beyond two weeks (Dejean et al., 2012; Thomsen et al., 2012; Piaggio et al., 2014; Barnes et al., 2014; Williams et al., 2018). However, signal persistence is dependent on the substrate, as soil samples can recover vertebrate signals greater than 2 months old (Andersen et al., 2012; Leempoel et al., 2020).

For airborne systems, in addition to known abiotic and biotic factors that encourage degradation, sources of eDNA may also become temporarily or permanently unavailable through particle settling on alternative surfaces to the capture substrate, such as the ground. The process of particle settling and potential resuspension by wind currents could thus prolong the detectable signal (Roger et al., 2022). Dry airborne particle deposition is influenced by factors such as higher wind speeds, which can allow large particles to remain suspended for longer periods and promote long distance transport (Aluko & Noll, 2006; Mandal et al. 2025). Moreover, wet deposition processes such as the “wash out” of particles from the atmosphere to the surface can occur during periods of rainfall (Seijo et al. 2025). In open-air terrestrial settings, eDNA persistence may differ from estimates obtained in an enclosed space, considering the spatial heterogeneity introduced by air flow patterns and other environmental conditions such as temperature and rainfall (Piaggio et al. 2025).

Airborne metabarcoding studies investigating community assemblages suggest that detections reflect recent biological activity. For instance, plant communities experienced compositional turnover within 24 hours of sampling (Lin et al. 2025). For vertebrate detections, distinct patterns were recorded between two-week sampling intervals (Johnson et al., 2023). Airborne signals are likely to persist beyond 12 hours, as no differences were observed in day versus night collected samples (Lynggaard et al., 2024). However, diurnal variation has been previously observed in detections of highly mobile bats, appearing to reflect extremely fine-scale temporal patterns in species activity (Garrett et al., 2023b). Daily patterns reflected in airborne eDNA are also consistent with evidence of landscape use and activity recovered from camera traps (Polling et al. 2024). An important consideration is that airborne eDNA persistence may differ by taxa, due to a variety of factors such as biomass, the state of the eDNA being shed, and activity patterns such as behaviour and mobility, which will affect the way eDNA is released into the air, where is it dispersed, and how much is being shed in different locations (Lynggaard et al., 2024).

The persistence of airborne eDNA signals has been previously investigated in an enclosed space by tracking detections associated with rare bat capture events (Garrett et al. 2026). Bat species of interest were commonly detected only on the night they were present in the room, or they were intermittently or consistently detected across several days following removal. Overall, the findings suggested a short temporal window of airborne eDNA persistence, up to 72 hours (3 days) post-removal, within which detections primarily reflect very recent biological activity, providing evidence of recent area use by the target. Based on these findings, Garrett et al. (2026) postulated that detecting relatively rare species would require increased sampling effort, with sampling needing to occur during or shortly after species presence. In contrast, they anticipate that abundant or common taxa may remain detectable in airborne samples for longer durations as more DNA is being shed. These align with findings for individuals versus groups in other eDNA mediums, such as from terrestrial swab samples (Piaggio et al. 2025) and in freshwater samples (Williams et al. 2018), whereby the signal left by a single individual degraded within days, whereas a group of the target species left a signal that persisted beyond the sampling timeframe of several weeks to months.

Here, we test temporal airborne eDNA persistence in an open-air natural setting, using the Bennett’s wallaby *Notamacropus rufogriseus* (Desmarest, 1817), a pest species in parts of its range (Latham et al., 2020), as a targeted case study with potential management application. Kroos et al. (2026) previously evaluated airborne eDNA detection of *N. rufogriseus* in captivity, but sampled continuously while the source population remained present, leaving open the question of how long an airborne eDNA signal persists once a source is removed, which has direct implications for practical deployment in operational surveillance. Moreover, Kroos et al. (2026) sampled across a distance gradient of 10-1000m using a captive source population of 27 individuals, and found that detection probability declined sharply with distance from the source. Their detection models indicated that reliably detecting eDNA from a single individual would require substantially greater sampling effort at distance (e.g., at 10 metres distance, an estimated 79 active airborne field samples would be required to detect a single individual compared to 3 field samples to detect the captive population of 27 individuals). Currently, single-individual airborne eDNA detectability at distance remains unvalidated empirically in open-air settings; however, this represents an important consideration since operational deployment is likely to occur in contexts involving individuals or low-density populations.

We also revisit two further questions initially raised by Kroos et al. (2026). Firstly, they found that passive airborne samplers without fans performed significantly worse than active (fan-powered) samplers paired side-by-side at close ranges to the source, but detection rates between the two methods converged at greater distances, with no significant difference observed at 1 km away from the source. Repeating this comparison under different source and distance conditions provides a further opportunity to clarify how collection method interacts with distance from the source. Secondly, Kroos et al. (2026) assessed several environmental variables as potential predictors of detection probability, but found that only sampler orientation relative to wind direction emerged as significant, albeit with wide confidence intervals, while variables such as wind speed showed no clear effect. Given this limited and uncertain set of findings, we examine the influence of environmental conditions on airborne eDNA detection in a new context.

In the present study, we quantify the temporal persistence of the airborne eDNA signal following source removal in an open-air, natural setting, and test the sensitivity of airborne eDNA detection from a single known-source individual at distance. We utilize passive (no-fan) and active (fan-powered) samplers paired side-by-side, to assess the impact of sampler type on detection probability, and measure environmental variables throughout the experiment to assess the impact of environmental factors. We captured airborne eDNA from a single Bennett’s wallaby carcass, deployed in an area where wallabies are otherwise not present, specifically, a nature reserve outside of the invasive range of wallabies. We tested several hypotheses: (1) airborne eDNA would be detectable shortly after carcass deployment and persist after carcass removal; (2) detection probability would significantly decline with distance; (3) detection probability would significantly differ by sampling method (active versus passive sampling); and (4) environmental variables, such as wind conditions, would significantly influence detection probability. Controlled conditions involving a single known source of wallaby DNA and an immobile target allowed us to assess airborne eDNA accumulation, transport, and persistence without including complex parameters such as animal movement.

## 2 METHODS

### 2.1 Field sampling

One adult female Bennett’s wallaby carcass weighing 13.8 kg and measuring 0.62 metres in length from the head to the beginning of the tail was sourced from roadkill in Central Otago, South Island, New Zealand, and stored in a freezer at –20 °C for six months prior to the experiment. Approximately 1.5 hours before deployment, it was removed from the freezer and left at ambient temperature. The carcass was placed on a flat, grassy surface within a wire mesh cage to prevent scavenging by animals (Figure 1A).

**Figure 1.**
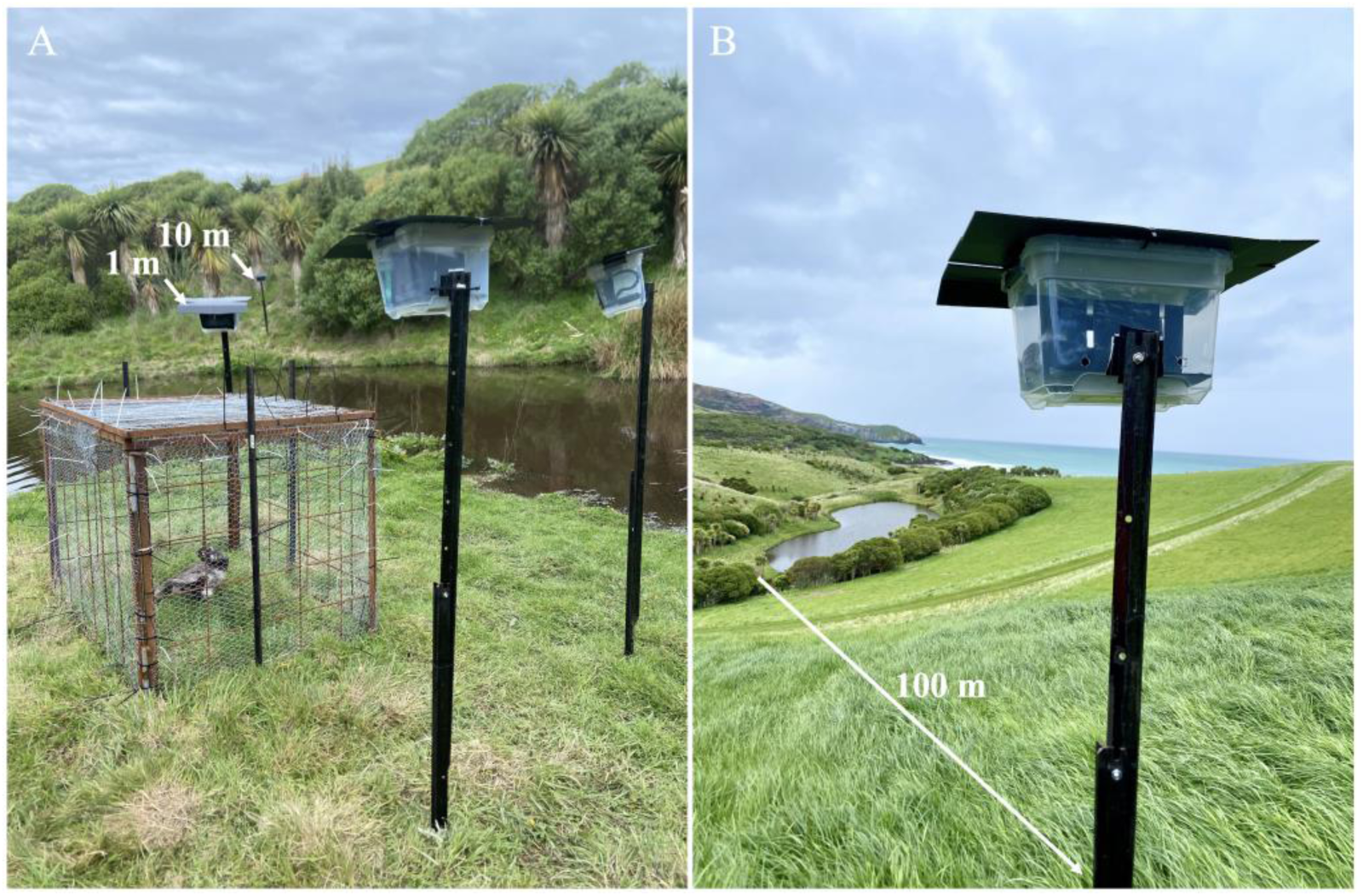
Airborne eDNA samplers utilizing both active and passive collection approaches side-by side deployed at 1.5 metre heights at various distances in relation to a single known source of wallaby DNA, at the Otago Peninsula, Dunedin, New Zealand. (A) Airborne eDNA samplers deployed at distances of 1 metre and 10 metres from a known source of wallaby DNA. (B) An airborne eDNA sampler deployed at a distance of 100 metres from a known source of wallaby DNA.

The carcass was deployed in an area that is known to be free of wallabies, specifically, the Otago Peninsula Eco Restoration Alliance, hereafter known as “the OPERA”, at Harington Point, Dunedin, New Zealand (45°48’19”S, 170°44’21”E). The OPERA sits at the tip of the Otago Peninsula, a coastal headland mainly composed of pasture and shrubs, experiencing a relatively dry, temperate maritime climate with cool temperatures (5–18 °C) and frequent, strong southwesterly and northeasterly winds (Macara, 2015). While Bennett’s wallabies have been reported elsewhere in the Otago region, they have never been recorded on or near the Otago Peninsula. Public sightings of wallabies, or their scat/footprints, have not been recorded within 20 kilometres of our sampling site (Wallaby Information System for the National Wallaby Eradication Programme, accessed 10 March 2026). As an added precaution, air samples were collected prior to the introduction of the wallaby carcass to serve as negative site controls.

For airborne particle capture we used the active and passive collection approaches outlined in Kroos et al. (2026). Briefly, a fibreglass media roll (Filters Direct, New Zealand), rated G4 under EN 779:2012 and suitable for capturing coarse particles larger than 10 µm, was pulled into layers, each ∼10 mm thick, and cut to 160 mm squares in a dedicated eDNA pre-PCR laboratory. Prior to deployment, each side of the material was exposed to UV light for 30 minutes, then the filters were stored in individual, sterile Ziploc bags. At each site, one active and one passive sampler were positioned side-by-side, housed in 6.4L plastic containers mounted 1.5 metres above ground level by attaching them to steel Y-posts (Figure 1). A rectangular cutout (160 mm by 60 mm) on the front face of each container allowed unobstructed airflow directly onto the filters. The active approach used a battery-powered computer fan to draw additional air through the filter, and the passive approach relied solely on ambient air movement.

Airborne eDNA sampling was conducted at nine locations, with three sites at each of three distances measured from the edge of the cage holding the wallaby carcass: 1 metre, 10 metres, and 100 metres (Figures 1, 2). Site selection and sampler orientation were guided by the prevailing southwesterly and northeasterly wind directions on the Otago Peninsula (Table S1). Samplers were exposed to air for durations ranging from 21 hours and 23 minutes to 25 hours and 49 minutes (Tables S2-S16). After sampling, filters were removed, wearing gloves and using sterile forceps, folded in half with the exposed side inward, and stored individually in sterile Ziploc bags. Samples were kept on ice in a cooling box for up to six hours before transfer to a –20 °C freezer.

**Figure 2.**
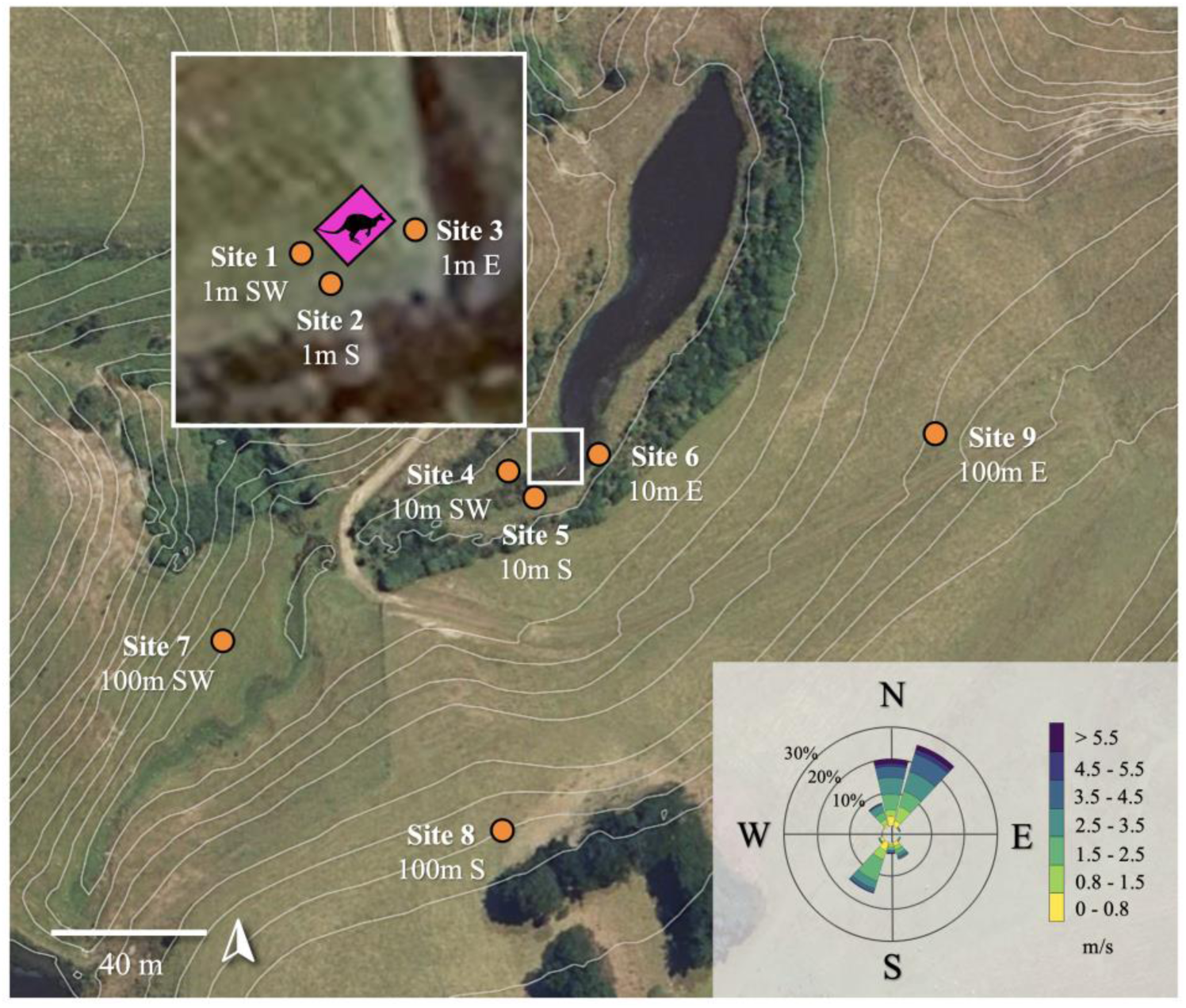
Airborne eDNA sampling sites (orange) at the Otago Peninsula, Dunedin, New Zealand, in relation to a single known source of wallaby DNA (pink), (45°48’19”S, 170°44’21”E). Average wind conditions for the entire sampling period are included, such as the frequency (%) of directions that winds were travelling from, and wind speeds measured in metres per second (m/s). Site topography is illustrated by contour intervals every 2 metres, generated using drone-based LIDAR.

The experiment was conducted over a six-week period, from 29 October -12 December 2024. Air sampling was conducted prior to carcass deployment, with sampling occurring at each of the nine sites, yielding 18 samples in total to serve as negative site controls. For 11 days that the carcass was present on site, air sampling occurred at six time points. After the carcass was removed, air sampling occurred at eight time points over the following 32 days to monitor the strength, direction, and persistence of the airborne eDNA signal in the absence of the wallaby DNA source.

Air samples were collected from nine sites, yielding 18 samples in total, per time point. Exceptions to this protocol occurred following carcass removal: at 11 days post-removal, sampling was restricted to sites at 1 metre and 10 metre distances, yielding 12 samples in total at this time point, and at 18, 25, and 32 days post-removal, sampling was restricted to sites at 1 metre distances, yielding six samples per time point. A total of 108 air samples (54 active, 54 passive) were collected during the period in which the carcass was present. After the removal of the carcass, a total of 102 air samples (51 active, 51 passive) were collected.

To minimise the risk of DNA contamination between sites and sampling periods, the following precautions were implemented. The first replicate of non-control samples were deployed prior to the wallaby carcass being handled and brought to the site. The samples were deployed by 2:59 pm, and the carcass was deployed at 4:45 pm on 30/10/24. Prior to being brought to the site, the carcass was stored in a freezer at the New Zealand Marine Studies Centre in Portobello, which was approximately 11.4 kilometres away. The carcass was stored inside two layers of clean plastic bags during transport to and from the site, and handled wearing sterile gloves. When the carcass was removed from the site, the next replicate of samples were deployed over 24 hours later. The carcass was removed at 9:58 am on 10/11/24, and the next replicate of samples began to be deployed at 11:58 am on 11/11/24. After being removed from the site, the carcass was transported to a nearby farm and put into an underground disposal pit.

Each day, sites were visited in order from the most distant (100 metres) to the closest (1 metre) to the wallaby carcass, beginning at site 9, and following sites in descending numerical order ending at site 1. Sampler components were disassembled on a sterilised plastic fold-out table, while wearing gloves and disinfected by soaking in 10% bleach (v/v) for 10 minutes, followed by a rinse with distilled water after each sampling period. Components were then reassembled for the next use. Gloves were changed between handling uncleaned and cleaned equipment, as well as between every sampling site. Field personnel also wore new, clean clothing each day.

Local meteorological conditions including wind direction, wind speed, rainfall, temperature, and humidity were obtained at hourly intervals using a single digital weather station (IC-XC0432, Digitech) facing due south, which was held 1.5 metres above ground, positioned 1 metre distance from the wallaby carcass. We collected meteorological data to account for the influence of environmental conditions such as rainfall, wind speed, and wind direction on airborne samples (Johnson et al., 2023).

### 2.2 DNA Extraction

DNA extraction was carried out in a dedicated eDNA pre-PCR laboratory, with several measures to reduce risk of contamination, including a unidirectional workflow, the wearing of hairnet, coveralls, two layers of medical gloves, allocated footwear, decontamination of surfaces and equipment using 10% bleach, and exclusive use of filter tips. DNA was extracted using the DNeasy Blood & Tissue Kit (QIAGEN, USA) following the manufacturer’s instructions, with the following modifications. Using sterile scissors, filters were cut into 60 mm squares taken from the centre, then each piece was halved and placed into separate 2 mL Eppendorf LoBind tubes using sterile forceps. After digestion, the corresponding halves of each air filter sample were recombined onto a single spin column. Filters were digested overnight at 56 °C with constant agitation at 1000 rpm. Each digestion tube contained 720 µl ATL buffer and 80 µl Proteinase K. Modified volumes of AL buffer and ethanol (600 µl each per sample) were used during extraction. DNA was eluted twice with 40 µl AE buffer per elution, each incubated at 37 °C for 15 minutes, resulting in a total elution volume of 80 µl. In addition to extracting field samples, negative filter controls were included, composed of sterilised air filter materials exposed to UV light for 30 minutes on both sides (n = 2). Additionally, one negative extraction control was included for every 23 field samples to detect any potential contamination in the extraction reagents.

### 2.3 qPCR Analysis

All qPCR reactions were set up in a dedicated eDNA pre-PCR laboratory separate from the eDNA extraction laboratory, following the same protocols to reduce contamination risk as outlined above. We used N.rufo primers and probe (Kroos et al., 2026) targeting the mitochondrial NADH dehydrogenase gene (MT-ND2 gene) of *N. rufogriseus*. This assay has multiple mismatches occurring in the forward, reverse, and probe regions to closely related/ co-occurring mammalian species in New Zealand as well as high sensitivity to detect the target, with a limit of detection (LOD) of 4.5 copies of target DNA per 1 µl.

Reactions were run on a QuantStudio 3.0 Real-time PCR instrument (Life Technologies) in 20 µl volumes; each containing 0.8 µl of the forward and reverse primers, and 0.2 µl of the probe (all 10 µM working stock),10 µl 2x SensiFAST probe mix (ThermoFisher Scientific), 6.2 µl nuclease-free water, and 2 µl template DNA. Thermocycling conditions were as follows: 95°C for 3 minutes, 95 °C for 15 seconds, annealing at 58 °C for 30 seconds and 50 cycles, followed by 72 °C for 20 seconds. Field-collected samples were analysed alongside multiple negative controls, including negative extraction controls, negative filter controls, and PCR negative controls (n = 3 per plate). A standard curve was also included on each plate, consisting of a 10-fold dilution series of known target DNA copy numbers ranging from 64,880 - 0.06488 copies per 2 µl reaction, with each standard concentration run in triplicate, using synthetic, double-stranded oligonucleotides of the *N. rufogriseus* MT-ND2 gene, consisting of the 106 bp PCR product flanked by additional 20 bp sequences on either end (total length 146 bp) inserted into a research-grade DNA plasmid (pUC57 vector) (Kroos et al., 2026).

A field sample was considered positive for *N. rufogriseus* DNA if it met the following criteria: 1) the amplification curve crossed the threshold for detection within 40 cycles, 2) uniform curve morphology, and 3) no detection observed in negative control samples (Hunter et al., 2017; Klymus et al., 2020). A field sample was considered positive for wallaby DNA if one, two, or three technical replicates produced a positive detection.

### 2.4 Statistical analyses

All statistical analyses were conducted in R version 4.5.1 (R Core Team, 2025). Detection probabilities were calculated for each distance category as the proportion of positive samples (i.e., samples had at least one positive technical replicate). These probabilities were then used to estimate the number of field replicates required to achieve a 95% probability of at least one positive detection at each distance using a binomial probability model.

We applied generalised linear models (GLMs) with a binomial distribution to model detection outcomes as binary variables (positive = 1, negative = 0). These models allowed us to assess the fixed effects of distance and environmental conditions on detection probability. Fixed environmental variables were; **Temperature**: The average air temperature during the sampling period, measured in degrees Celsius. **Humidity**: The average relative humidity percentage during the sampling period. **Wind Speed**: The average wind speed during the sampling period, measured in metres per second. **Wind Direction**: an average of the direction from which the wind was blowing from, expressed in degrees from true north (0° indicating wind from the north) for the sampling period. Compass directions were converted to degrees, from degrees to radians, and transformed from a linear variable to *sin* and *cos* components. **Rainfall**: The total amount of precipitation that occurred during the sampling period, measured in millimetres. **Site Direction**: The compass direction that the sampler was facing at each site, expressed in degrees from true north, with data transformed the same as for wind direction. Backward model selection was performed with the *dredge* function from the MuMIn package v1.46.0 (Bartoń, 2022), starting from the most comprehensive model including all predictors. Models were ranked by Akaike Information Criterion corrected for small sample size (AICc), and the best-fitting models were chosen based on the lowest AICc values. Wald tests (Pr(>|z|)) were used to determine the significance of fixed effects. Models were considered significant if *p* < 0.05 and 95% bootstrap confidence intervals excluded zero. We also conducted model averaging including models within 4 AICc units of the top model.

A correlation matrix was generated to examine relationships among predictor variables. To address potential multicollinearity among environmental predictors, we performed principal component analysis (PCA) on scaled variables including wind speed, wind direction, temperature, humidity, and site direction. The resulting principal components (PCs), were then used as predictors in a logistic regression GLM to examine their collective influence on detection probability. Models with increasing numbers of PCs (1 to 3) were compared, and model averaging was applied to synthesise results across models.

To compare detection outcomes between active and passive airborne samplers collected concurrently at the same sites and days, we conducted an exact binomial test on active detections, to find out if the detection rate using active significantly differs from the detection rate of passive samplers.

Nonparametric Kruskal-Wallis rank-sum tests were conducted to assess differences in cycle quantification (Cq) values, and DNA quantity (DNA copies/µl) in relation to distance from the source and time elapsed since carcass introduction/ time elapsed since carcass removal. Where significant differences were found, post-hoc pairwise comparisons were performed using Dunn’s test with Bonferroni correction.

To investigate potential changes in DNA copy number over time since wallaby carcass addition in the experiment, we applied segmented regression models to the DNA quantity data (copies/µl) using the *segmented* package v2.1-4 (Muggeo, 2008). The time variable was expressed in minutes elapsed since the carcass was added. This approach estimates a breakpoint at which the relationship between DNA quantity and time shifts. A Wilcoxon Rank Sum test was then applied to compare DNA quantity and Cq values from samples collected before versus after carcass removal.

## 3 RESULTS

Negative site controls collected prior to the introduction of the wallaby carcass, as well as all filter controls, extraction controls, and PCR controls, returned negative results for Bennett’s wallaby DNA.

Following carcass introduction to the site, 6 of 90 samples collected using active air sampling were positive, yielding an overall detection rate of 6.67%, with DNA copy numbers in all positive samples above the assay’s LOD. For 90 samples collected using passive air sampling, none returned a positive detection. An exact binomial test revealed a significant difference in detection rate using active versus passive sampling (p < 2.2 x 10⁻¹⁶). No positive detections occurred at sites that were visited immediately after sites that tested positive (minimum separation = 6 site visits, occurring 7 days post-deployment at site 2, then 8 days post-deployment at site 5) (Figure 3), suggesting carryover contamination between consecutively visited sites was unlikely.

**Figure 3.**
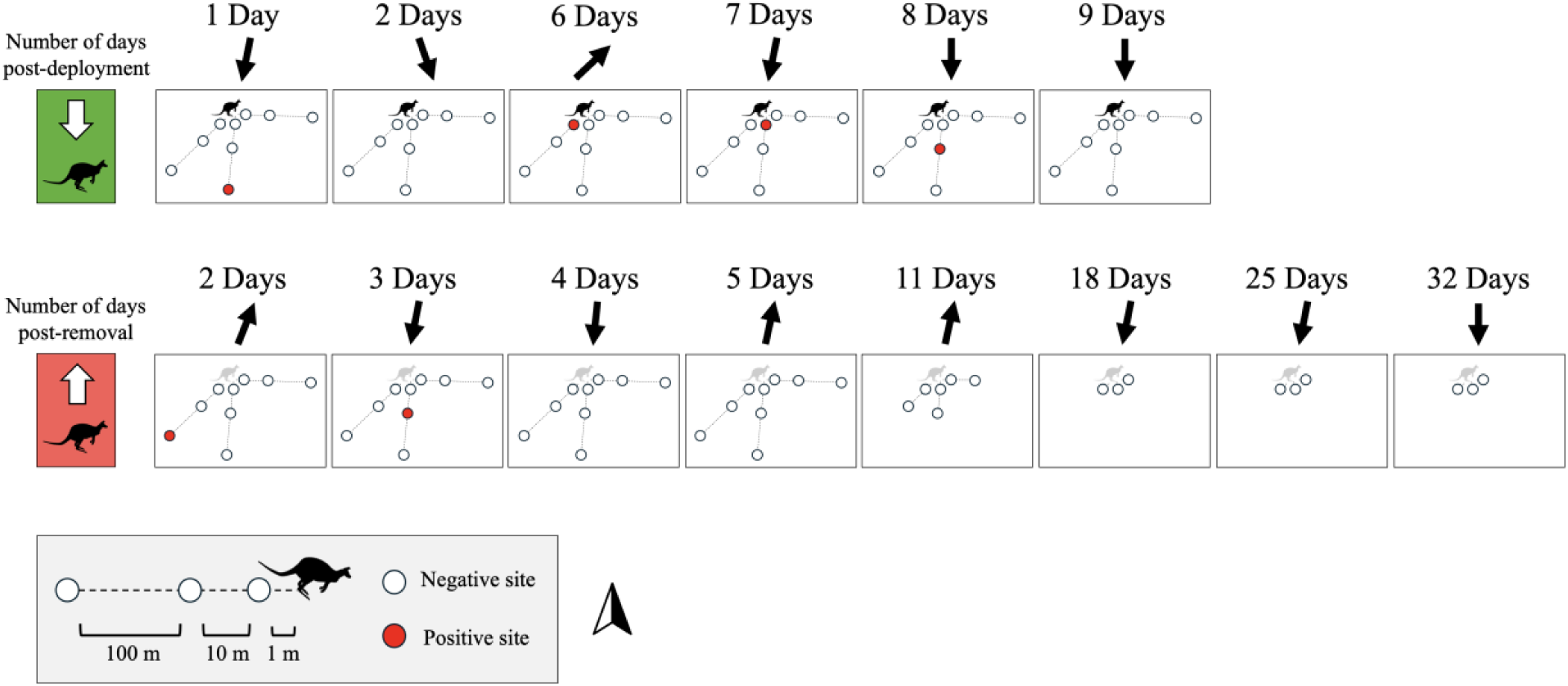
Positive detections from airborne eDNA samples across days since the wallaby DNA was deployed and days since the wallaby DNA was removed, as well as the origin of positive detections in relation to the source of wallaby DNA. Black arrows indicate the prevailing direction the wind was blowing from during the specified sampling period.

Wallaby DNA was detected from airborne samples collected 18 hours after wallaby carcass deployment, and this first positive detection originated 100 metres away. The carcass remained detectable up to 8 days after its initial deployment into the environment while it was still present, with the quantity of DNA (copies/µl) from samples collected during carcass presence being the highest in the sample collected 6 days after deployment, 1 metre distance away (Table S17). Following carcass removal, positive detections originated from a site 100 metres away, 2 days after removal, and 10 metres away, 3 days after removal. Positive detections ceased after 3 days post-removal (Figure 3).

The DNA concentration (copies/µL) recovered from positive samples were analysed in relation to the addition and removal of the wallaby DNA source (Figure 4). The estimated breakpoint at which a shift in DNA quantity occurred was after 5.9 days of the carcass being present on site; however, model uncertainty was high (SE ≈ 4,158), the overall model fit was very low (R² = 0.016), and neither pre- nor post-breakpoint slopes were statistically significant. Additionally, no significant temporal effects were detected for DNA quantity or Cq values using Wilcoxon rank sum tests (quantity: p = 0.681; Cq: p = 0.692).

**Figure 4.**
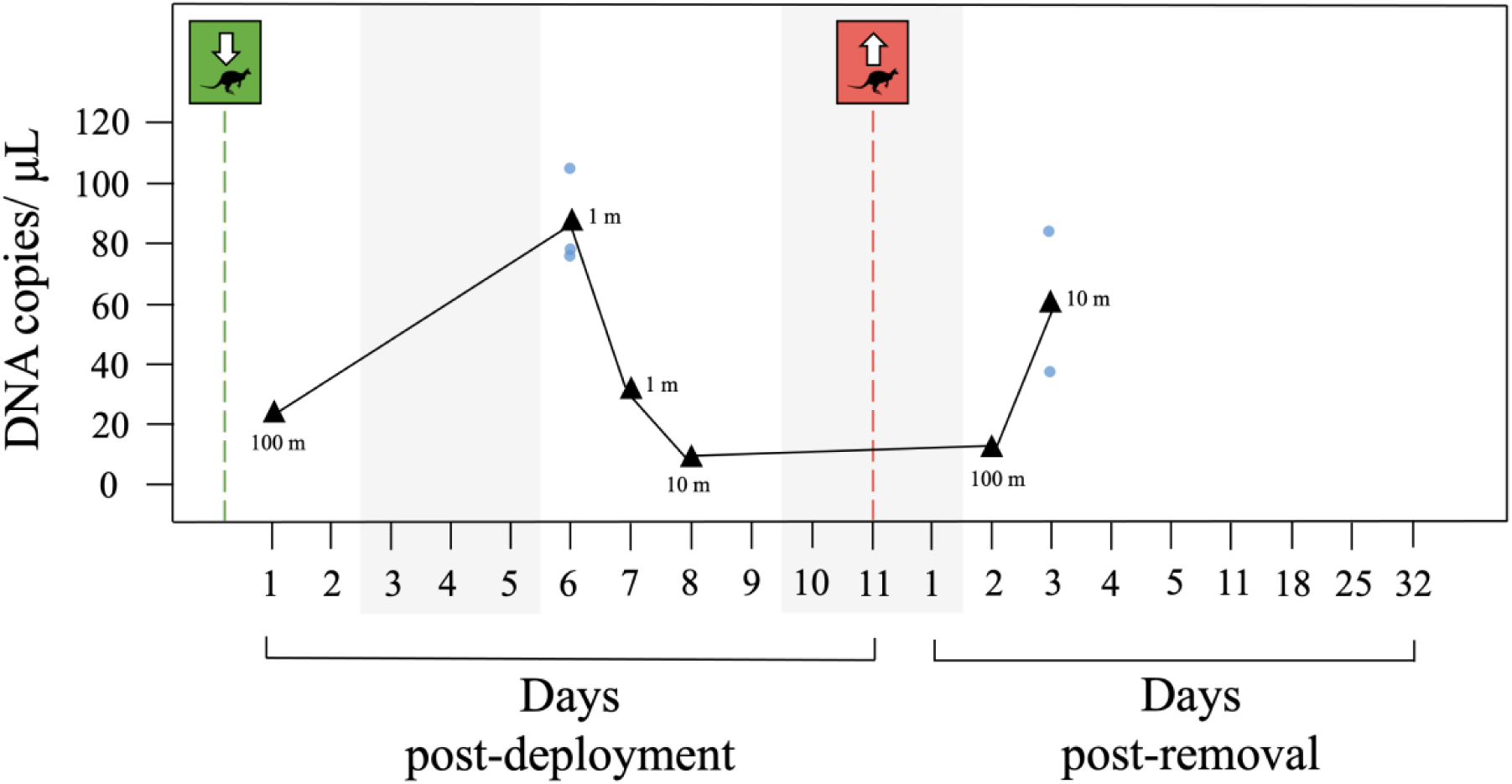
Average wallaby DNA quantity from positive samples (black triangle) collected at distances of 1 metre, 10 metres, or 100 metres from a known source of wallaby DNA. Grey represents days that sampling did not occur.

At each sampling distance, 2 of 30 samples collected using active sampling were positive. A Kruskal-Wallis test of Cq values across distance groups (1 metre, 10 metres, and 100 metres) detected a significant overall effect of Cq values increasing with distance (p = 0.0498). However, post-hoc pairwise comparisons using Dunn’s tests with Bonferroni correction revealed no significant differences between any distance groups (1 m–10 m: p = 0.094; 1 m–100 m: p = 0.173; 10 m–100 m: p = 1.0). Consistent with this, a Kruskal–Wallis test comparing estimated DNA quantity (copies/µl) among distance groups showed no significant differences (p = 0.233).

We then used a binomial probability model to estimate the future sampling effort required for reliable detection. Assuming a consistent detection probability of 6.67% at each distance category, at least 45 field replicates would be required per distance (1 metre, 10 metres, and 100 metres) to achieve at least one positive detection of a single wallaby with 95% confidence.

During the experimental period, total recorded rainfall was 16.6 mm, with a maximum of 6.6 mm falling within a single 24-hour period. Temperatures ranged from 4.7°C to 18.8°C. The highest recorded wind speed was 7.6 m/s, with an average hourly wind speed of 1.78 m/s. The prevailing wind direction was northeasterly (Fig 2). Assessing the influence of environmental factors on airborne detection rates, the best-fitting model included the variable *Site_direction_cos*, representing site orientation relative to wind direction. However, its association with detection probability was not statistically significant (p = 0.128). Several models performed similarly to this model, as such, model averaging was performed to account for uncertainty. Model averaging revealed no significant effects for any environmental variable on detection rates, including rainfall, humidity, temperature, wind speed, wind direction, distance, or site orientation.

A correlation matrix of environmental variables revealed several correlations including strong positive correlations between temperature and humidity (r = 0.58), wind speed and wind direction (r = 0.85), and temperature and wind direction (r = 0.61), all with p < 0.001. To address multicollinearity, principal component analysis (PCA) was applied to the scaled environmental predictors, with the first three principal components (PCs) explaining 81% of the total variance. Model-averaged coefficients from the PCA-based models revealed no statistically significant predictors of detection (p > 0.05). Overall, neither individual environmental variables nor their combined structure in a PCA significantly explained variation in detection probability in this dataset.

However, we observed a strong correlation between prevailing wind direction measured over each 24-hour sampling period and the spatial distribution of sites that tested positive for wallaby airborne eDNA (Figure 3). For four of the six positive detections, winds predominantly originated from the northeast or north, and positive detections occurred at sites where sampling devices were positioned to intercept air travelling from the north (sites 8, 5, and 2, Table S1).

Overall, following deployment of a wallaby carcass into an open-air natural environment where wallabies are otherwise absent, wallaby DNA was detected from airborne sampling 18 hours later at a distance of 100 metres from the introduction site. Positive detections persisted up to three days after carcass removal, after which no further wallaby DNA was detected. Active air sampling using battery-powered fans significantly improved detection rates compared to passive sampling. While no environmental variables were significantly correlated with detection rates, we observed an association between detection events and the alignment between the airborne sampler’s orientation and the prevailing wind direction.

## 4 DISCUSSION

### 4.1 Temporal persistence

In support of our hypothesis, airborne eDNA sampling yielded a positive detection shortly after deploying a DNA source, and the positive signal persisted following removal of the source. We found that within 18 hours of introduction to a previously wallaby-free site, wallaby DNA was detected 100 metres from the release point using airborne sampling. This rapid detection suggests that airborne signals can disperse quickly after deposition, traveling hundreds of metres and likely beyond. Moreover, these results indicate our targeted assay shows strong potential as a sensitive early-detection tool for this species, at low target densities such as a single individual.

Positive detections occurred twice after the carcass was removed, and ceased after three days post-removal, suggesting a short persistence of the airborne DNA signal in an open-air environment. Possible explanations for this trend are that particles carrying most of the airborne eDNA signal are quickly transported away from the local area, or, remain suspended for only brief periods before settling onto surfaces, where they may become unavailable to samplers, or degrade. Notably, our findings align with persistence estimates derived from terrestrial surface swabs collecting eDNA, whereby a detection window of four days was observed after source animals (mice) were removed from an open-air arena, before DNA concentrations likely fell below detectable levels (Piaggio et al. 2025). This similarity suggests that, in open-air terrestrial environments, airborne eDNA and eDNA deposited onto surfaces may have similarly short detection windows following source removal, potentially owing to comparable exposure to environmental factors that facilitate DNA degradation.

Our findings are very consistent with previous observations in a partially enclosed space that airborne eDNA signals are lost within a few days of the source being removed (Garrett et al. 2026). This concordance between studies suggests that airborne eDNA detections could be generally indicative of recent species presence or activity, regardless of whether sampling occurs in enclosed or open-air environments. However, important differences exist between these systems. In enclosed and partially enclosed settings, air movement is more constrained, whereas open-air environments are subject to greater transport of eDNA particles away from the sampled area through factors such as heightened wind activity, as well as the impacts of rainfall, which may reduce airborne particles available to be captured (Macher et al., 2023; Seijo et al. 2025), as well as greater exposure to sunlight and temperature fluctuations, which may heighten degradation rates of airborne particles (e.g., Strickler et al., 2015). Consequently, airborne sampler placement and sampling effort may play a more critical role in open systems to minimise false negative results, whereby the target is present in the environment but is missed.

Analysis of DNA quantity over time revealed an initial accumulation of wallaby DNA in airborne samples, followed by a subsequent decline. Under controlled conditions, eDNA tends to exponentially increase through time when the source remains present, and decline over time following the removal of the source (Williams et al. 2018; Piaggio et al., 2025). However, we observed a decline in wallaby DNA before source removal, suggesting that environmental degradation or dispersal mechanisms may have started to reduce airborne DNA availability prior to the disappearance of the physical source.

Nonparametric tests showed no significant correlation between time since carcass placement or removal and either DNA quantity (copies/µL) or Cq values, which we suspect is likely as a result of our limited dataset, with high temporal variability and noise, and few positive detections. Although Wilcoxon tests were applied to assess changes in DNA quantity and Cq over time, the number of observations may have been insufficient.

Our airborne detections were infrequent, and spatially and temporally inconsistent rather than continuous. Similarly, Garrett et al. (2026) reported intermittent detections of target bat species, with signals from rare taxa becoming detectable at later time points despite the animals not being present or detected via eDNA in the intervening period. They suggested that the most likely explanation for these observations was the resuspension of eDNA following disturbance within the enclosed space they sampled. In our study, we observed a secondary “spike” in detections after the carcass was removed, occurring several days after a non-detection event on the last day that the carcass was present on site (9 days post-deployment, Figure 4). This pattern may also be attributed to the resuspension of particles into the air. Such resuspension could have resulted from physical disturbance due to our own movement in the area, increased wind activity, an increase in dryness following a period of moderate rainfall, or a combination of these factors.

We observed that the average wind speed increased for the two sampling periods directly after carcass removal (from 0.91 m/s on the final day of carcass presence on site, to 1.36, then 2.0 m/s, on the days following removal). Predominantly northerly and north-northeasterly winds, with gusts of up to 7.4 m/s were recorded on the day of carcass removal, and may also account for the detection of wallaby eDNA at sampling site 7, situated 100 metres away, 2 days post-removal of the carcass. The orientation of this site, and the orientation of the sampling device relative to the carcass is consistent with the prevailing wind direction during this period (Figure 3; Table S1). We also observed that the highest rainfall during a single sampling period (6.6 mm within 4 hours) coincided with the sampling time point that was 9 days post-deployment of the carcass, which had no positive detections, despite positive detections the previous 3 days. Positive detections resumed two days after this period, during which the carcass was removed, with no rainfall in the intervening time.

Our infrequent detection pattern also likely reflects the low concentration of vertebrate eDNA in outdoor air, consistent with previous findings (e.g., Lynggaard et al., 2024), and is likely to be even rarer when originating from a single individual source. Moreover, insects have a known role in the transport and transfer of genetic material (Calvignac-Spencer et al. 2013; Massey et al. 2022; Fernandes et al. 2024). As such, insect activity may have contributed to sporadic detection events, particularly at further distances from the source. This mechanism is plausible given that insects, primarily sandflies and blow flies, were observed on the carcass, directly on the filter material, and resting on the exterior surfaces of the airborne samplers. Larger scavengers were excluded by the metal cage, but insects and smaller scavengers could still act as important vectors for terrestrial vertebrate airborne eDNA, particularly from a deceased source.

### 4.2 Distance effect

Active airborne sampling using fan power successfully detected wallaby eDNA from a single individual source at distances of 1, 10, and 100 metres in an open-air, natural setting. Although dispersal of airborne eDNA over hundreds of metres from the nearest known source has been reported previously in captive settings (Jager et al. 2026) and in natural environments (Frère et al. 2024), we demonstrate here that airborne eDNA from one individual can rapidly disperse over distances of at least 100 metres, which will be informative for management applications. However, our results did not support our hypothesis that detection probability significantly decreases with increasing distance from the DNA source. Typically, higher detection rates are observed closer to the source animal (e.g., Lynggaard et al. 2024; Newton et al., 2024; Kroos et al. 2026) because of rapid dilution as distance from the source increases, and since larger, heavier airborne particles often travel shorter distances (Tournayre et al. 2025). Although no statistically significant relationship was found between distance and DNA quantity, the highest DNA yields occurred at the closest sampling distance (1 metre), which is consistent with our expectations.

We suspect that detection rates at the 1 metre sites would have been higher relative to the 10 metre and 100 metre sites if our airborne samplers had been positioned closer to the ground. The carcass was lying flat on the ground, while samplers were deployed at a height of approximately 1.5 metres, likely reducing the probability of intercepting airborne particles in close proximity to the source. At greater distances, wind may facilitate vertical transport of particles, increasing the likelihood of interception at the sampler height. The detection patterns we observed may also reflect the detection of eDNA from a single individual rather than the more abundant signal emitted by a group of the species of interest; consequently, detection rates may be more sporadic, even at very close ranges, thus any detection patterns with distance did not emerge. Another consideration is that site topography varied considerably across some of the sampling locations. Two sites located 100 metres from the source were positioned on large hills, 12-16 metres above the elevation of the carcass, whereas the rest of the sites were at the same height as the carcass (Figure 2). Differences in elevation may have enhanced or reduced the capacity of these elevated sites to intercept airborne particles, and is a factor worth exploring in future studies considering that natural landscapes vary widely in topography.

### 4.3 Collection method

Our results support our hypothesis that detection probability significantly differs by sampling collection method when comparing active, fan-powered sampling versus passive, no-fan sampling. Passive airborne samplers failed entirely to detect wallaby eDNA in these experiments, despite previous evidence of successful detection from passive samplers at distances of up to 1 kilometre from a captive wallaby enclosure (Kroos et al. 2026). This discrepancy is likely attributable to differences in target abundance. In the present study, we aimed to detect an individual signal, rather than the substantially higher eDNA signal expected from dense captive populations. Active samplers may have outperformed passive samplers because they processed a greater volume of air, or alternatively, the action of the fan created a difference in pressure which was more efficient at capturing particles on the filter substrate. Although passive versus active sampler air volumes weren’t compared directly, the active samplers are known to process an air volume of 66.49 m^3^/hour when a filter is attached (Kroos et al. 2026). Since the samplers were deployed side by side, differences were not attributable to location, direction, or differential exposure. Taken together, these findings suggest that passive airborne samplers may have limited utility for detecting extremely rare targets.

### 4.4 Environmental variables

Our results do not support our hypothesis that environmental variables, such as wind, significantly influence detection probability. We found that temperature, humidity, wind speed, rainfall and wind direction did not exhibit strong or consistent effects on airborne eDNA detection probability from our models. Overall, our inability to resolve significant environmental influences likely reflects limitations in sample size that hinder the ability to disentangle the influence of environmental variability. However, sampler orientation emerged repeatedly as a variable of interest. Although not statistically significant, a positive association between sampler orientation and detection suggests that alignment with prevailing winds, mediated by local topography and wind conditions, influences how airborne eDNA accumulates or disperses. Consistent with these findings, we did observe correlation between prevailing wind direction and the orientation of samplers that were positive. Kroos et al. (2026) also identified sampler orientation as a strong predictor of airborne detectability. Based on our findings, we would recommend future studies incorporate a sampler that can capture airborne particles originating from all directions.

### 4.5 Limitations and Caveats

An important limitation of our study is that we used a carcass rather than a live individual as a source of DNA. While we did find that a single wallaby carcass shed detectable airborne eDNA, we acknowledge that sampling a carcass, which represents a static DNA source, does not fully reflect the spatial and temporal dynamics of DNA shedding and dispersal associated with live animal movement, as well as animal behaviour, which is an important aspect to interpreting airborne signals (Johnson et al., 2023). In natural systems, live animals contribute to airborne eDNA through various behaviours such as breathing, sneezing, scratching, and rubbing against surfaces, all of which can increase DNA shedding. At the same time, their mobility can disperse this DNA more widely, potentially reducing concentrations at any single location. These dynamic factors are likely to play a significant role in shaping airborne eDNA detection patterns in natural environments.

Moreover, carcasses are likely to shed eDNA differently than live animals. Aquatic eDNA research has shown that fish carcasses can release detectable eDNA for extended periods as tissues break down and cellular material is released (Merkes et al. 2014). Tillotson et al. (2018) found dead fish to have a higher DNA shedding rate than live fish, which likely has important implications for this study, though it is unknown whether similar processes occur in airborne systems. In addition, a carcass may attract greater insect activity than a live animal; therefore, insect-mediated transport of DNA may have been more likely to have contributed to positive detections in this study. The carcass we used had also been stored long term at −20 °C prior to deployment and was allowed to thaw for only 1.5 hours before being placed in the environment. Consequently, the deposition and accumulation of airborne eDNA may have differed from that of a naturally deceased animal, particularly during the thawing out process.

We also recognise that our estimates of airborne persistence may not be directly transferable to other contexts, and may differ depending on the taxa, environment, and airborne sampling method. Our estimates of persistence are derived from a relatively large-bodied, ground-dwelling mammal covered in hair, potentially enhancing the rate of detection, as there is some evidence to suggest that taxa with larger biomass are positively correlated with higher read counts, suggesting that larger vertebrates release more DNA into the air (Jager et al., 2026). Moreover, we used airborne filters that are rated to capture larger particle sizes (greater than 10 μM), which are currently not a widely utilized choice for filters in airborne eDNA studies. Although it is not yet well understood, the cellular form and particle size of airborne eDNA may influence the deposition of eDNA and its persistence (Lin et al. 2025; Garrett et al. 2026). Finally, since we used a method that could capture air originating only from one direction, we expect that the window of persistence may be extended further if our samplers were able to capture wind from all directions.

### 4.6 Implications for Management

Promisingly, our estimates of the number of airborne eDNA samples needed to detect a single wallaby in an open air setting were substantially lower than those reported by Kroos et al. (2026), who calculated that 477 samples would be required to achieve 95% confidence of at least one detection at distances up to 100 metres. Based on our observed detection rate of 6.67% across all distances, only 45 replicate samples would be needed to achieve at least one positive detection of an individual up to 100 metres away, reducing sampling effort more than tenfold. Differences may have arisen because our estimates are based on empirical detection rates from a single individual, whereas Kroos et al. (2026) derived their estimate from modelled detection probabilities based on a larger population and extrapolated these to the expected detectability of an individual. These results suggest that, from a management perspective, detecting rare targets may require fewer airborne eDNA samples than previously thought to achieve the necessary sensitivity.

Based on the timeframe over which airborne wallaby DNA remained detectable, we can infer management-relevant detection windows (Williams et al. 2018). Available evidence suggests that if a wallaby were removed through management intervention (e.g., a control operation), detection of airborne wallaby eDNA in the weeks thereafter would most likely indicate the presence of a new individual. These temporal thresholds may require adjustment where multiple individuals are suspected to be present, as residual eDNA could be greater and persist for longer (Garrett et al. 2026). Physical removal of wallabies killed by control operations, as well as incorporating a buffer period of at least one week following a control operation, would therefore substantially reduce the likelihood of false-positive detections attributed to remnant DNA. Moreover, our findings suggest that for future management scenarios, sampler replication and sampling duration will need to account for short-term persistence of airborne signals in open-air environments.

Overall, we found that airborne eDNA could detect a single vertebrate individual in an open-air, natural setting at distances of up to 100 metres, and for up to three days following removal of the DNA source, providing evidence that eDNA persistence in air is short-term. This understanding of temporal persistence of airborne eDNA is critical for optimising field deployment strategies. Detection rates were low, likely reflecting the sporadic and rare occurrence of target DNA in air samples from an individual source, and may be mediated by environmental conditions such as wind direction. Active air collection using battery-powered fans substantially improved the probability of detecting airborne eDNA from an individual source, compared to passive air collection without the use of fans. Our results highlight the capacity of airborne eDNA to detect individual organisms in outdoor environments, while underscoring the need for robust sampling approaches and replication to capture rare and transient DNA signals.

## Supplementary information

**Table S1.**
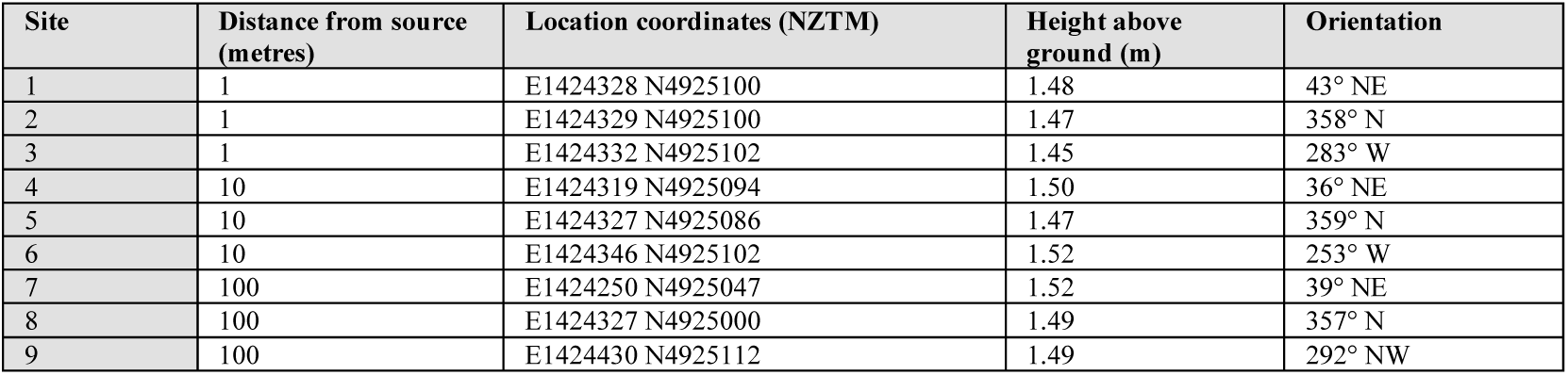
Otago Peninsula air sampling site details, including site ID, distance from the source of wallaby DNA (wallaby carcass), location coordinates, height above ground (in metres), and sampler orientation.

**Table S2.**
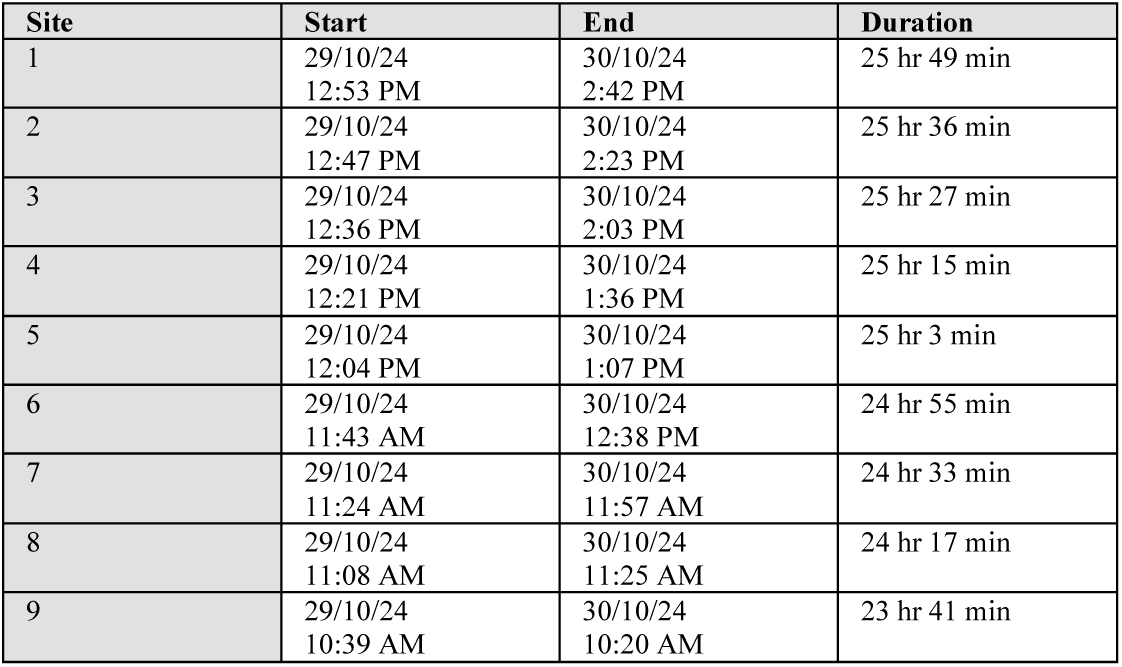
Otago Peninsula air sampling details for negative control samples, which were collected prior to wallaby deployment on 30/11/24.

**Table S3.**
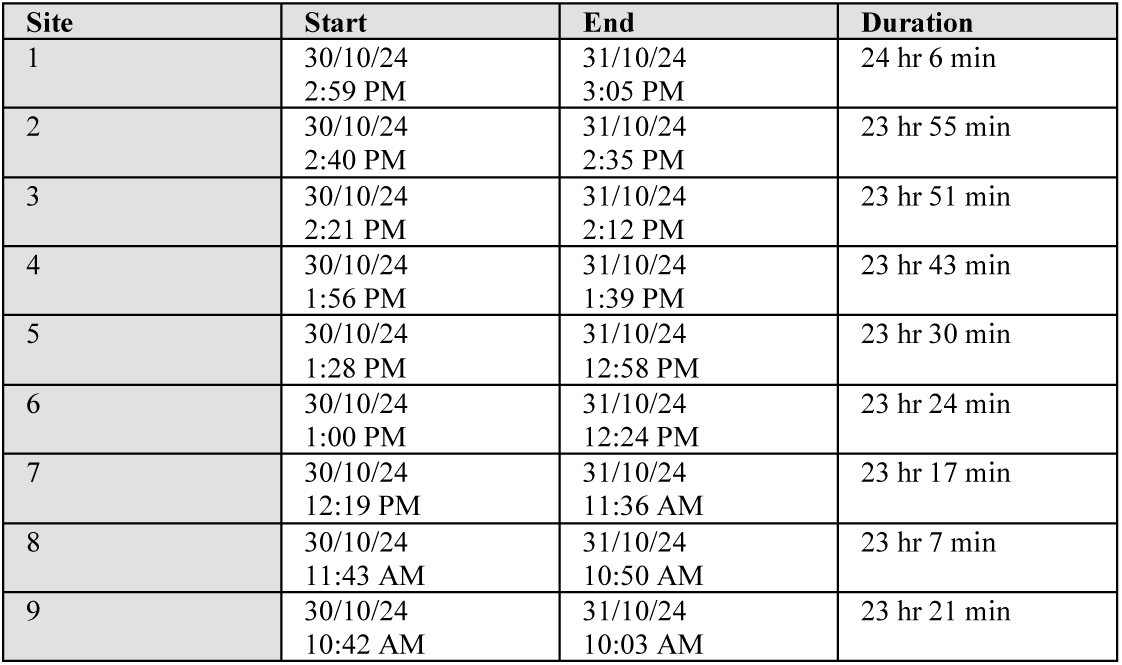
Otago Peninsula air sampling details for samples collected 1 day post-deployment on 31/10/24.

**Table S4.**
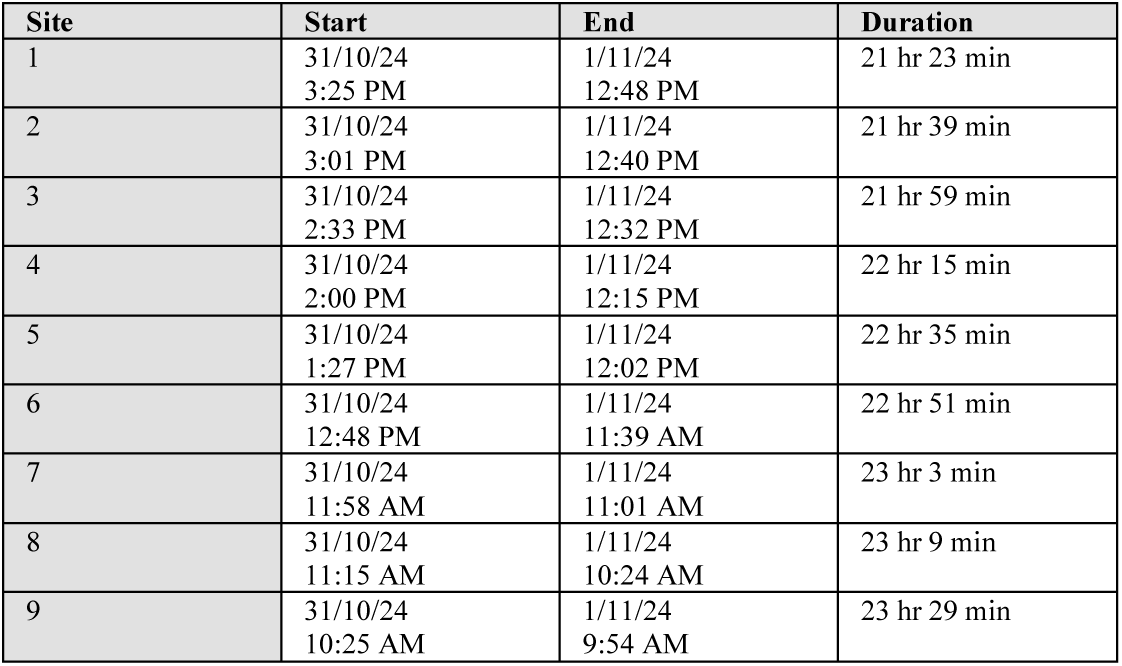
Otago Peninsula air sampling details for samples collected 2 days post-deployment on 1/11/24.

**Table S5.**
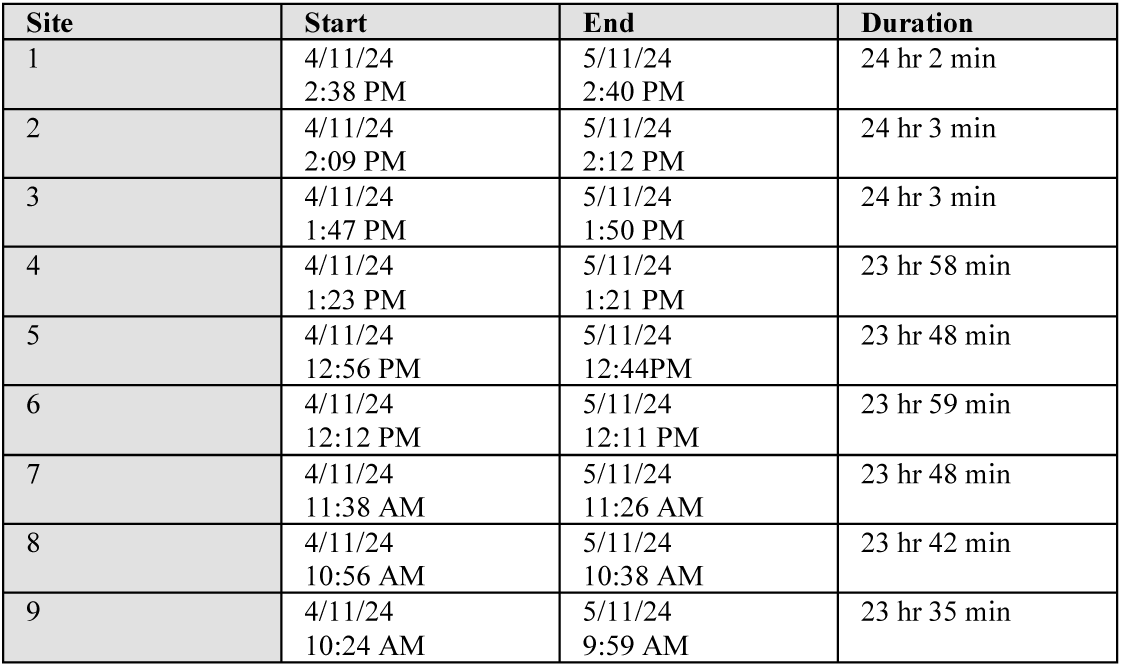
Otago Peninsula air sampling details for samples collected 6 days post-deployment on 5/11/24.

**Table S6.**
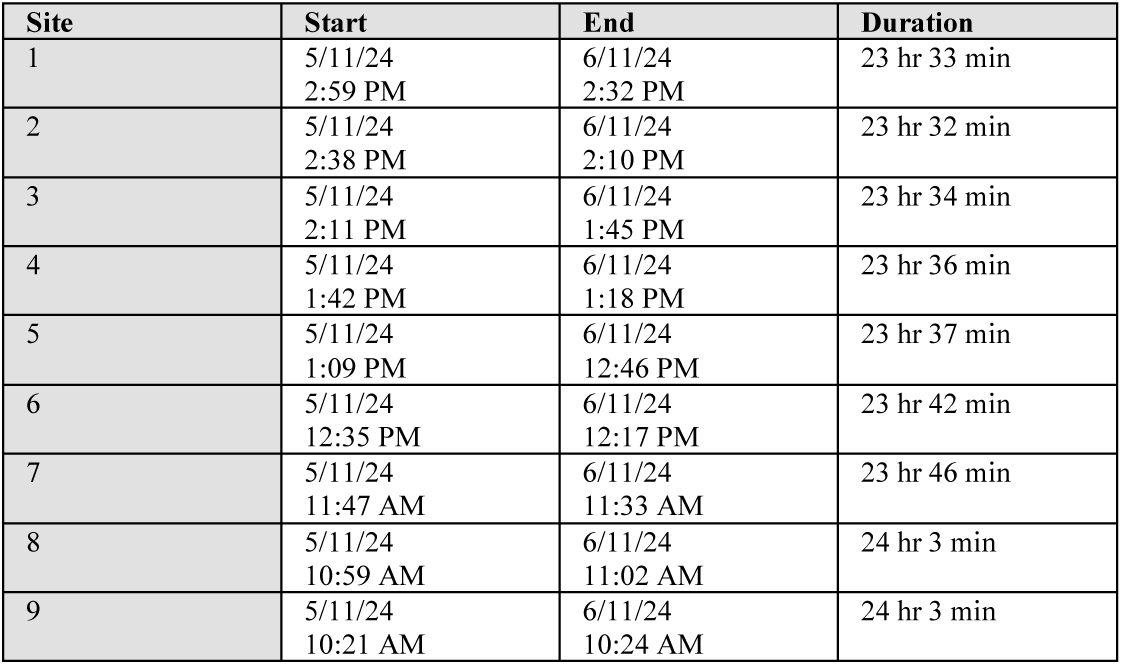
Otago Peninsula air sampling details for samples collected 7 days post-deployment on 6/11/24.

**Table S7.**
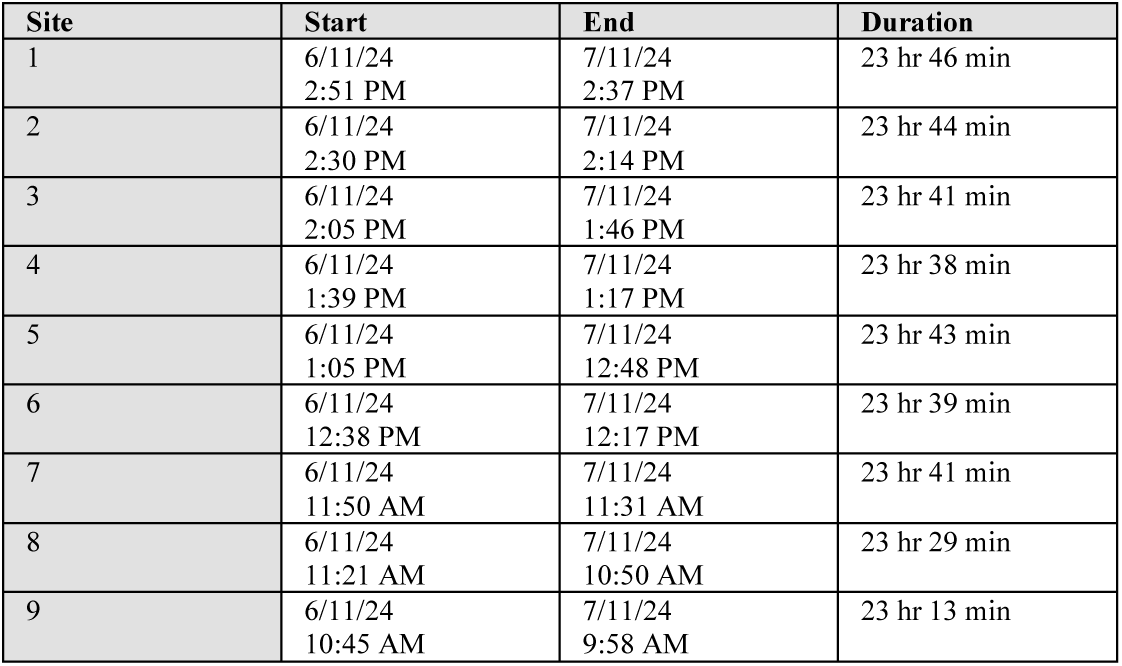
Otago Peninsula air sampling details for samples collected 8 days post-deployment on 7/11/24.

**Table S8.**
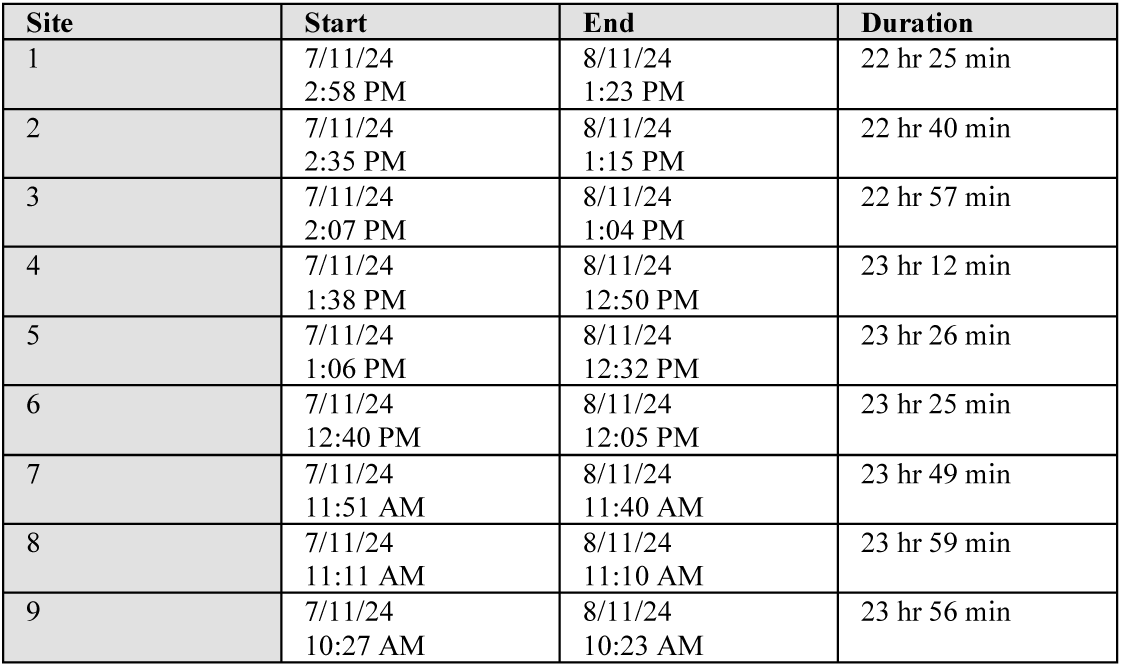
Otago Peninsula air sampling details for samples collected 9 days post-deployment on 8/11/24.

**Table S9.**
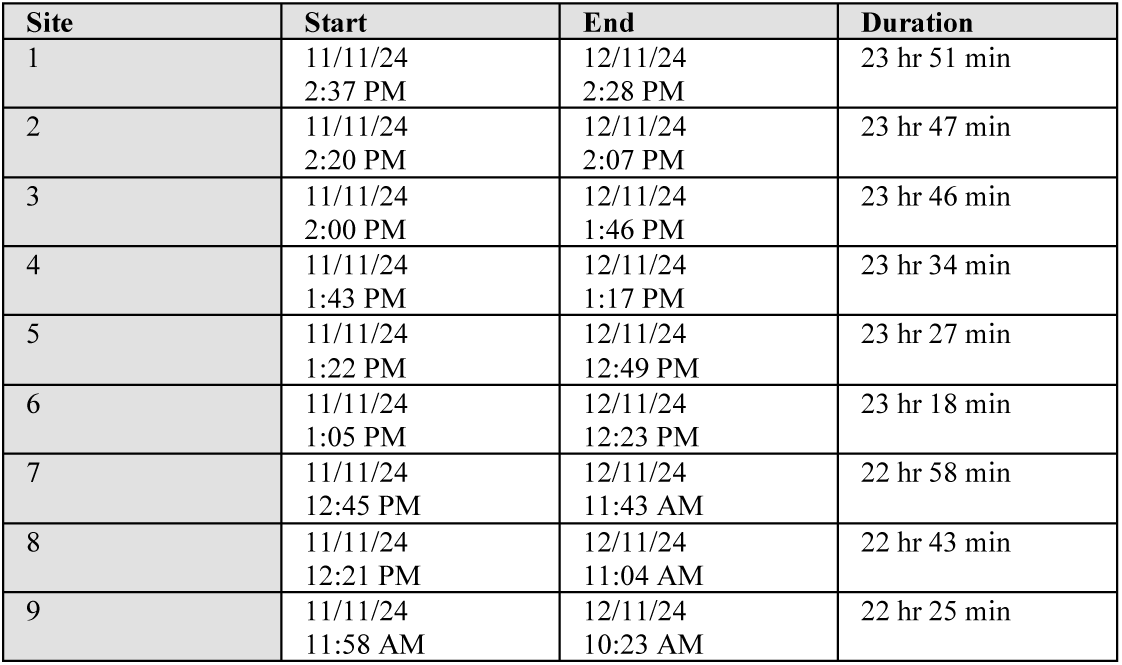
Otago Peninsula air sampling details for samples collected 2 days post-removal on 12/11/24.

**Table S10.**
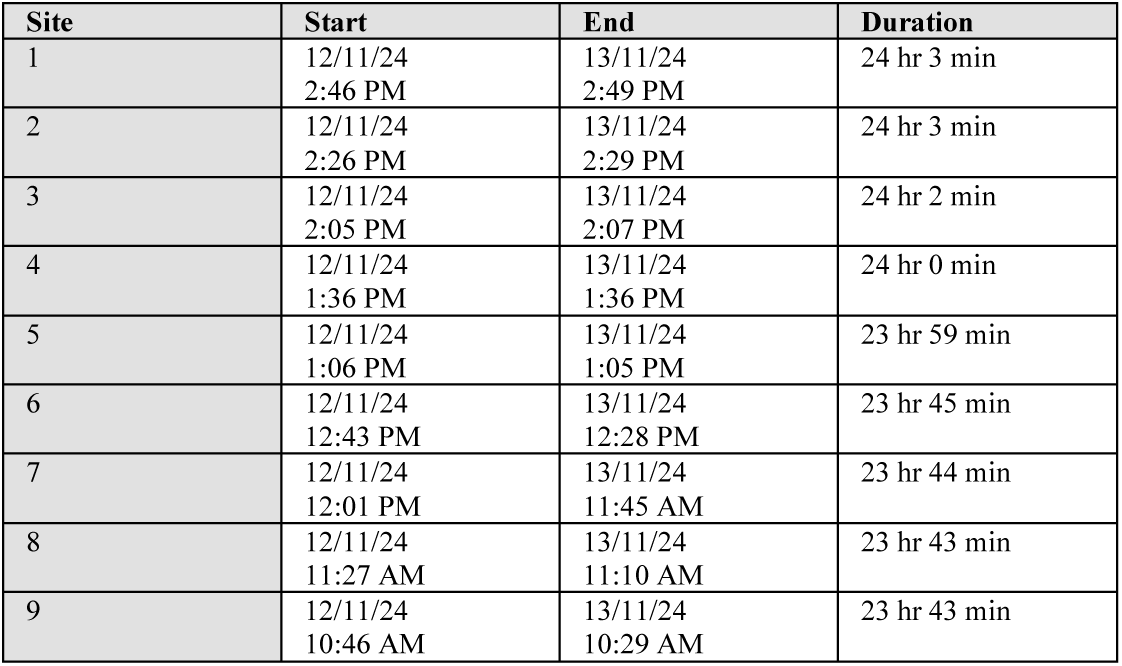
Otago Peninsula air sampling details for samples collected 3 days post-removal on 13/11/24.

**Table S11.**
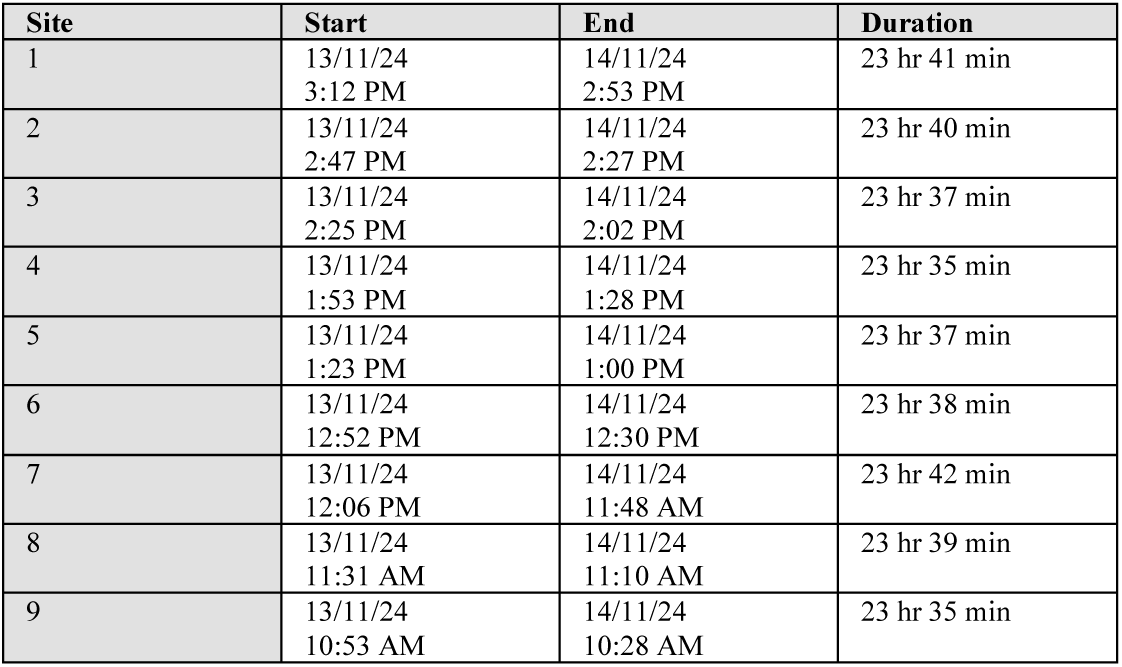
Otago Peninsula air sampling details for samples collected 4 days post-removal on 14/11/24.

**Table S12.**
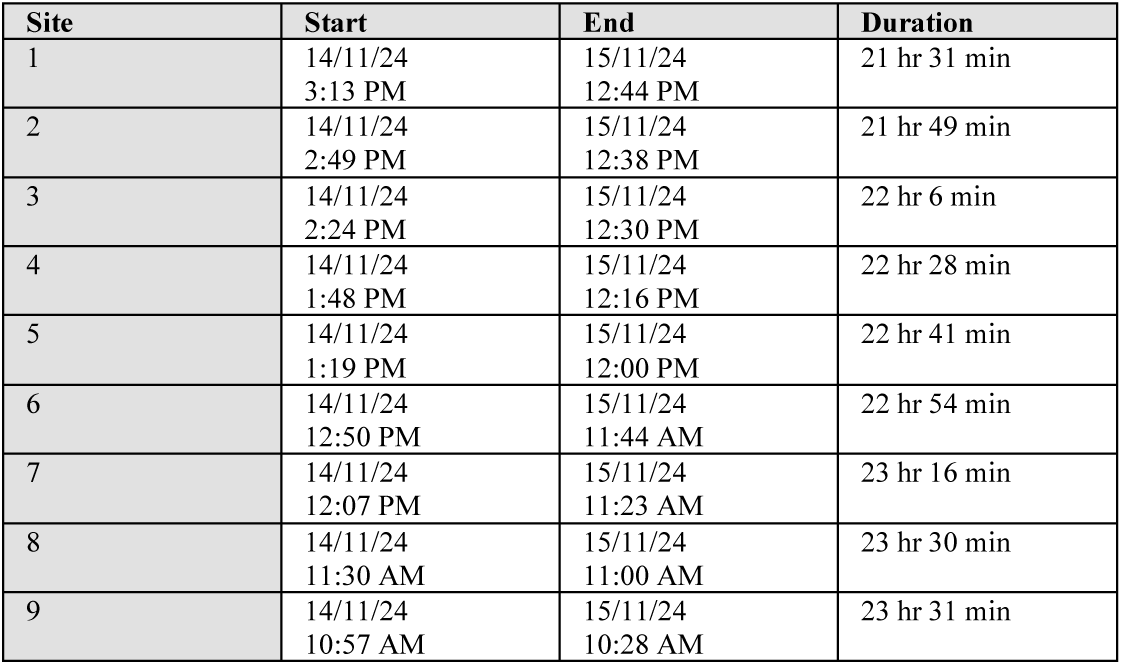
Otago Peninsula air sampling details for samples collected 5 days post-removal on 15/11/24.

**Table S13.**
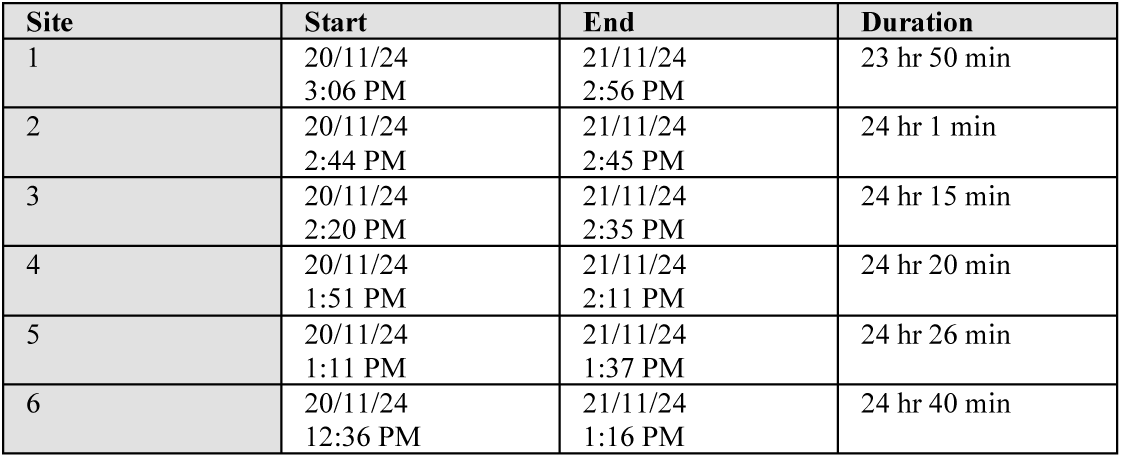
Otago Peninsula air sampling details for samples collected 11 days post-removal on 21/11/24.

**Table S14.**
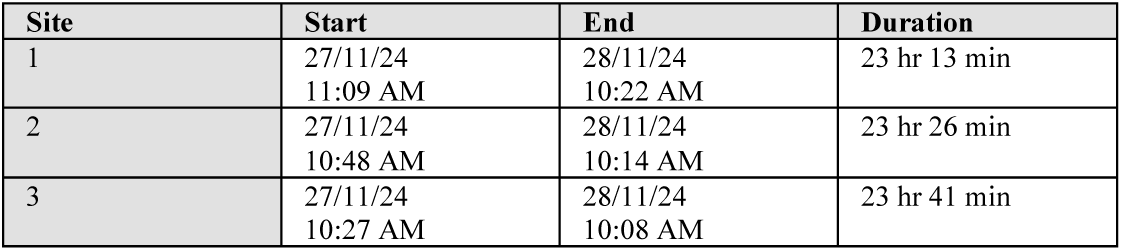
Otago Peninsula air sampling details for samples collected 18 days post-removal on 28/11/24.

**Table S15.**
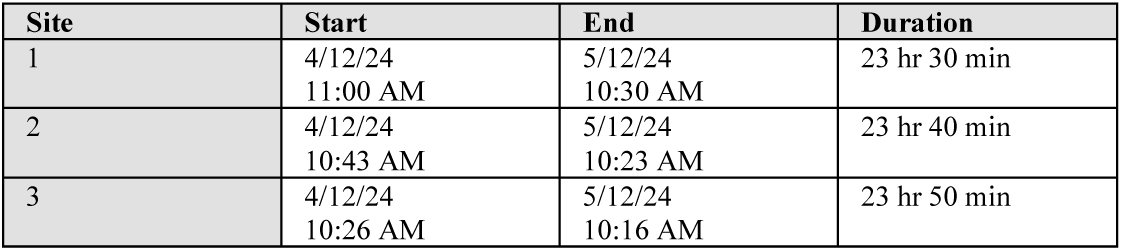
Otago Peninsula air sampling details for samples collected 25 days post-removal on 5/12/24.

**Table S16.**
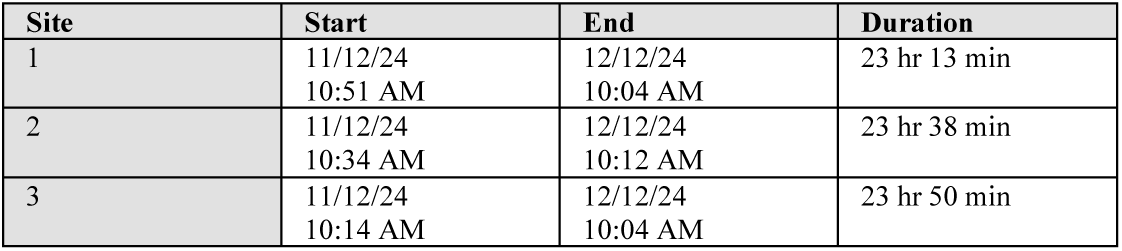
Otago Peninsula air sampling details for samples collected 32 days post-removal on 12/12/24.

**Table S17.**
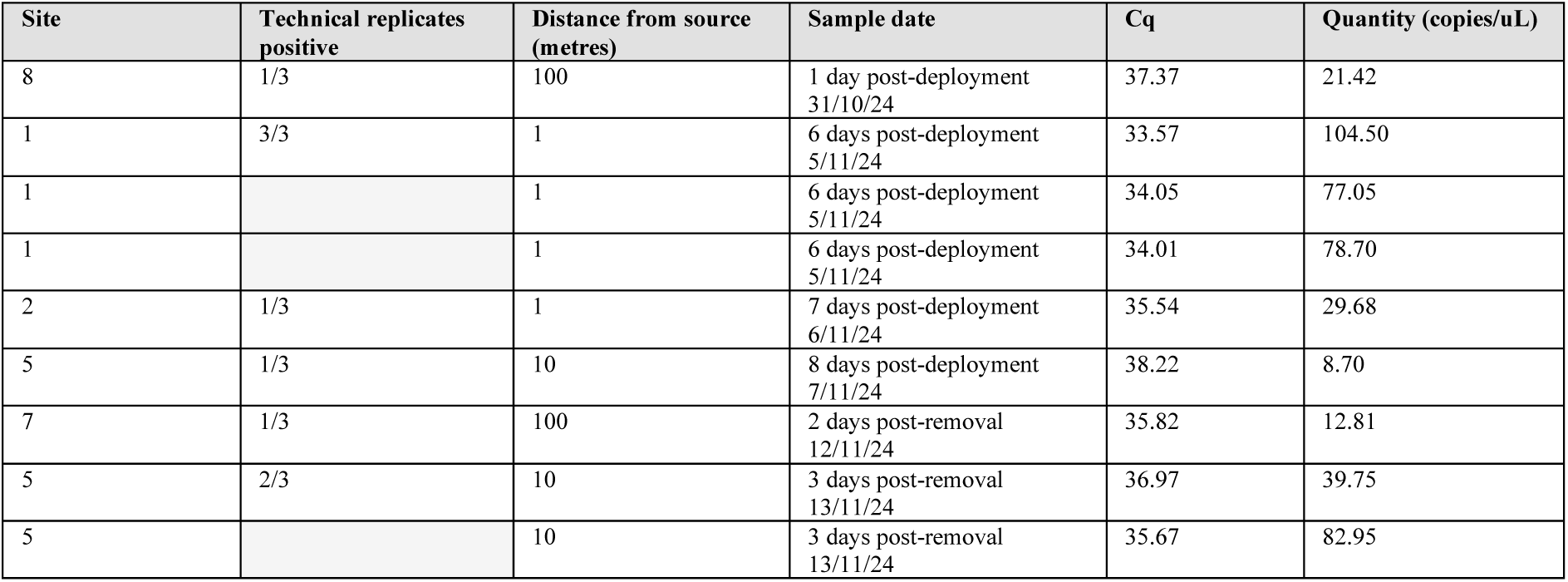
Otago Peninsula positive air sampling results including Cq value and DNA Quantity (copies/uL).

