## Supplementary information for "Temporal persistence of airborne environmental DNA in a natural open-air setting"

**Supplementary information- Temporal persistence of airborne eDNA**

**Tables S1-S17**

**Table S1. Otago Peninsula air sampling site details, including site ID, distance from the source of wallaby DNA (wallaby carcass), location coordinates, height above ground (in metres), and sampler orientation.**

| **Site** | **Distance from source (metres)** | **Location coordinates (NZTM)** | **Height above ground (m)** | **Orientation** |
| --- | --- | --- | --- | --- |
| 1 | 1 | E1424328 N4925100 | 1.48 | 43° NE |
| 2 | 1 | E1424329 N4925100 | 1.47 | 358° N |
| 3 | 1 | E1424332 N4925102 | 1.45 | 283° W |
| 4 | 10 | E1424319 N4925094 | 1.50 | 36° NE |
| 5 | 10 | E1424327 N4925086 | 1.47 | 359° N |
| 6 | 10 | E1424346 N4925102 | 1.52 | 253° W |
| 7 | 100 | E1424250 N4925047 | 1.52 | 39° NE |
| 8 | 100 | E1424327 N4925000 | 1.49 | 357° N |
| 9 | 100 | E1424430 N4925112 | 1.49 | 292° NW |

**Table S2. Otago Peninsula air sampling details for negative control samples, which were collected prior to wallaby deployment on 30/11/24.**

| **Site** | **Start** | **End** | **Duration** |
| --- | --- | --- | --- |
| 1 | 29/10/24  12:53 PM | 30/10/24  2:42 PM | 25 hr 49 min |
| 2 | 29/10/24  12:47 PM | 30/10/24  2:23 PM | 25 hr 36 min |
| 3 | 29/10/24  12:36 PM | 30/10/24  2:03 PM | 25 hr 27 min |
| 4 | 29/10/24  12:21 PM | 30/10/24  1:36 PM | 25 hr 15 min |
| 5 | 29/10/24  12:04 PM | 30/10/24  1:07 PM | 25 hr 3 min |
| 6 | 29/10/24  11:43 AM | 30/10/24  12:38 PM | 24 hr 55 min |
| 7 | 29/10/24  11:24 AM | 30/10/24  11:57 AM | 24 hr 33 min |
| 8 | 29/10/24  11:08 AM | 30/10/24  11:25 AM | 24 hr 17 min |
| 9 | 29/10/24  10:39 AM | 30/10/24  10:20 AM | 23 hr 41 min |

**Table S3. Otago Peninsula air sampling details for samples collected 1 day post-deployment on 31/10/24.**

| **Site** | **Start** | **End** | **Duration** |
| --- | --- | --- | --- |
| 1 | 30/10/24  2:59 PM | 31/10/24  3:05 PM | 24 hr 6 min |
| 2 | 30/10/24  2:40 PM | 31/10/24  2:35 PM | 23 hr 55 min |
| 3 | 30/10/24  2:21 PM | 31/10/24  2:12 PM | 23 hr 51 min |
| 4 | 30/10/24  1:56 PM | 31/10/24  1:39 PM | 23 hr 43 min |
| 5 | 30/10/24  1:28 PM | 31/10/24  12:58 PM | 23 hr 30 min |
| 6 | 30/10/24  1:00 PM | 31/10/24  12:24 PM | 23 hr 24 min |
| 7 | 30/10/24  12:19 PM | 31/10/24  11:36 AM | 23 hr 17 min |
| 8 | 30/10/24  11:43 AM | 31/10/24  10:50 AM | 23 hr 7 min |
| 9 | 30/10/24  10:42 AM | 31/10/24  10:03 AM | 23 hr 21 min |

**Table S4. Otago Peninsula air sampling details for samples collected 2 days post-deployment on 1/11/24.**

| **Site** | **Start** | **End** | **Duration** |
| --- | --- | --- | --- |
| 1 | 31/10/24  3:25 PM | 1/11/24  12:48 PM | 21 hr 23 min |
| 2 | 31/10/24  3:01 PM | 1/11/24  12:40 PM | 21 hr 39 min |
| 3 | 31/10/24  2:33 PM | 1/11/24  12:32 PM | 21 hr 59 min |
| 4 | 31/10/24  2:00 PM | 1/11/24  12:15 PM | 22 hr 15 min |
| 5 | 31/10/24  1:27 PM | 1/11/24  12:02 PM | 22 hr 35 min |
| 6 | 31/10/24  12:48 PM | 1/11/24  11:39 AM | 22 hr 51 min |
| 7 | 31/10/24  11:58 AM | 1/11/24  11:01 AM | 23 hr 3 min |
| 8 | 31/10/24  11:15 AM | 1/11/24  10:24 AM | 23 hr 9 min |
| 9 | 31/10/24  10:25 AM | 1/11/24  9:54 AM | 23 hr 29 min |

**Table S5. Otago Peninsula air sampling details for samples collected 6 days post-deployment on 5/11/24.**

| **Site** | **Start** | **End** | **Duration** |
| --- | --- | --- | --- |
| 1 | 4/11/24  2:38 PM | 5/11/24  2:40 PM | 24 hr 2 min |
| 2 | 4/11/24  2:09 PM | 5/11/24  2:12 PM | 24 hr 3 min |
| 3 | 4/11/24  1:47 PM | 5/11/24  1:50 PM | 24 hr 3 min |
| 4 | 4/11/24  1:23 PM | 5/11/24  1:21 PM | 23 hr 58 min |
| 5 | 4/11/24  12:56 PM | 5/11/24  12:44PM | 23 hr 48 min |
| 6 | 4/11/24  12:12 PM | 5/11/24  12:11 PM | 23 hr 59 min |
| 7 | 4/11/24  11:38 AM | 5/11/24  11:26 AM | 23 hr 48 min |
| 8 | 4/11/24  10:56 AM | 5/11/24  10:38 AM | 23 hr 42 min |
| 9 | 4/11/24  10:24 AM | 5/11/24  9:59 AM | 23 hr 35 min |

**Table S6. Otago Peninsula air sampling details for samples collected 7 days post-deployment on 6/11/24.**

| **Site** | **Start** | **End** | **Duration** |
| --- | --- | --- | --- |
| 1 | 5/11/24  2:59 PM | 6/11/24  2:32 PM | 23 hr 33 min |
| 2 | 5/11/24  2:38 PM | 6/11/24  2:10 PM | 23 hr 32 min |
| 3 | 5/11/24  2:11 PM | 6/11/24  1:45 PM | 23 hr 34 min |
| 4 | 5/11/24  1:42 PM | 6/11/24  1:18 PM | 23 hr 36 min |
| 5 | 5/11/24  1:09 PM | 6/11/24  12:46 PM | 23 hr 37 min |
| 6 | 5/11/24  12:35 PM | 6/11/24  12:17 PM | 23 hr 42 min |
| 7 | 5/11/24  11:47 AM | 6/11/24  11:33 AM | 23 hr 46 min |
| 8 | 5/11/24  10:59 AM | 6/11/24  11:02 AM | 24 hr 3 min |
| 9 | 5/11/24  10:21 AM | 6/11/24  10:24 AM | 24 hr 3 min |

**Table S7. Otago Peninsula air sampling details for samples collected 8 days post-deployment on 7/11/24.**

| **Site** | **Start** | **End** | **Duration** |
| --- | --- | --- | --- |
| 1 | 6/11/24  2:51 PM | 7/11/24  2:37 PM | 23 hr 46 min |
| 2 | 6/11/24  2:30 PM | 7/11/24  2:14 PM | 23 hr 44 min |
| 3 | 6/11/24  2:05 PM | 7/11/24  1:46 PM | 23 hr 41 min |
| 4 | 6/11/24  1:39 PM | 7/11/24  1:17 PM | 23 hr 38 min |
| 5 | 6/11/24  1:05 PM | 7/11/24  12:48 PM | 23 hr 43 min |
| 6 | 6/11/24  12:38 PM | 7/11/24  12:17 PM | 23 hr 39 min |
| 7 | 6/11/24  11:50 AM | 7/11/24  11:31 AM | 23 hr 41 min |
| 8 | 6/11/24  11:21 AM | 7/11/24  10:50 AM | 23 hr 29 min |
| 9 | 6/11/24  10:45 AM | 7/11/24  9:58 AM | 23 hr 13 min |

**Table S8. Otago Peninsula air sampling details for samples collected 9 days post-deployment on 8/11/24.**

| **Site** | **Start** | **End** | **Duration** |
| --- | --- | --- | --- |
| 1 | 7/11/24  2:58 PM | 8/11/24  1:23 PM | 22 hr 25 min |
| 2 | 7/11/24  2:35 PM | 8/11/24  1:15 PM | 22 hr 40 min |
| 3 | 7/11/24  2:07 PM | 8/11/24  1:04 PM | 22 hr 57 min |
| 4 | 7/11/24  1:38 PM | 8/11/24  12:50 PM | 23 hr 12 min |
| 5 | 7/11/24  1:06 PM | 8/11/24  12:32 PM | 23 hr 26 min |
| 6 | 7/11/24  12:40 PM | 8/11/24  12:05 PM | 23 hr 25 min |
| 7 | 7/11/24  11:51 AM | 8/11/24  11:40 AM | 23 hr 49 min |
| 8 | 7/11/24  11:11 AM | 8/11/24  11:10 AM | 23 hr 59 min |
| 9 | 7/11/24  10:27 AM | 8/11/24  10:23 AM | 23 hr 56 min |

**Table S9. Otago Peninsula air sampling details for samples collected 2 days post-removal on 12/11/24.**

| **Site** | **Start** | **End** | **Duration** |
| --- | --- | --- | --- |
| 1 | 11/11/24  2:37 PM | 12/11/24  2:28 PM | 23 hr 51 min |
| 2 | 11/11/24  2:20 PM | 12/11/24  2:07 PM | 23 hr 47 min |
| 3 | 11/11/24  2:00 PM | 12/11/24  1:46 PM | 23 hr 46 min |
| 4 | 11/11/24  1:43 PM | 12/11/24  1:17 PM | 23 hr 34 min |
| 5 | 11/11/24  1:22 PM | 12/11/24  12:49 PM | 23 hr 27 min |
| 6 | 11/11/24  1:05 PM | 12/11/24  12:23 PM | 23 hr 18 min |
| 7 | 11/11/24  12:45 PM | 12/11/24  11:43 AM | 22 hr 58 min |
| 8 | 11/11/24  12:21 PM | 12/11/24  11:04 AM | 22 hr 43 min |
| 9 | 11/11/24  11:58 AM | 12/11/24  10:23 AM | 22 hr 25 min |

**Table S10. Otago Peninsula air sampling details for samples collected 3 days post-removal on 13/11/24.**

| **Site** | **Start** | **End** | **Duration** |
| --- | --- | --- | --- |
| 1 | 12/11/24  2:46 PM | 13/11/24  2:49 PM | 24 hr 3 min |
| 2 | 12/11/24  2:26 PM | 13/11/24  2:29 PM | 24 hr 3 min |
| 3 | 12/11/24  2:05 PM | 13/11/24  2:07 PM | 24 hr 2 min |
| 4 | 12/11/24  1:36 PM | 13/11/24  1:36 PM | 24 hr 0 min |
| 5 | 12/11/24  1:06 PM | 13/11/24  1:05 PM | 23 hr 59 min |
| 6 | 12/11/24  12:43 PM | 13/11/24  12:28 PM | 23 hr 45 min |
| 7 | 12/11/24  12:01 PM | 13/11/24  11:45 AM | 23 hr 44 min |
| 8 | 12/11/24  11:27 AM | 13/11/24  11:10 AM | 23 hr 43 min |
| 9 | 12/11/24  10:46 AM | 13/11/24  10:29 AM | 23 hr 43 min |

**Table S11. Otago Peninsula air sampling details for samples collected 4 days post-removal on 14/11/24.**

| **Site** | **Start** | **End** | **Duration** |
| --- | --- | --- | --- |
| 1 | 13/11/24  3:12 PM | 14/11/24  2:53 PM | 23 hr 41 min |
| 2 | 13/11/24  2:47 PM | 14/11/24  2:27 PM | 23 hr 40 min |
| 3 | 13/11/24  2:25 PM | 14/11/24  2:02 PM | 23 hr 37 min |
| 4 | 13/11/24  1:53 PM | 14/11/24  1:28 PM | 23 hr 35 min |
| 5 | 13/11/24  1:23 PM | 14/11/24  1:00 PM | 23 hr 37 min |
| 6 | 13/11/24  12:52 PM | 14/11/24  12:30 PM | 23 hr 38 min |
| 7 | 13/11/24  12:06 PM | 14/11/24  11:48 AM | 23 hr 42 min |
| 8 | 13/11/24  11:31 AM | 14/11/24  11:10 AM | 23 hr 39 min |
| 9 | 13/11/24  10:53 AM | 14/11/24  10:28 AM | 23 hr 35 min |

**Table S12. Otago Peninsula air sampling details for samples collected 5 days post-removal on 15/11/24.**

| **Site** | **Start** | **End** | **Duration** |
| --- | --- | --- | --- |
| 1 | 14/11/24  3:13 PM | 15/11/24  12:44 PM | 21 hr 31 min |
| 2 | 14/11/24  2:49 PM | 15/11/24  12:38 PM | 21 hr 49 min |
| 3 | 14/11/24  2:24 PM | 15/11/24  12:30 PM | 22 hr 6 min |
| 4 | 14/11/24  1:48 PM | 15/11/24  12:16 PM | 22 hr 28 min |
| 5 | 14/11/24  1:19 PM | 15/11/24  12:00 PM | 22 hr 41 min |
| 6 | 14/11/24  12:50 PM | 15/11/24  11:44 AM | 22 hr 54 min |
| 7 | 14/11/24  12:07 PM | 15/11/24  11:23 AM | 23 hr 16 min |
| 8 | 14/11/24  11:30 AM | 15/11/24  11:00 AM | 23 hr 30 min |
| 9 | 14/11/24  10:57 AM | 15/11/24  10:28 AM | 23 hr 31 min |

**Table S13. Otago Peninsula air sampling details for samples collected 11 days post-removal on 21/11/24.**

| **Site** | **Start** | **End** | **Duration** |
| --- | --- | --- | --- |
| 1 | 20/11/24  3:06 PM | 21/11/24  2:56 PM | 23 hr 50 min |
| 2 | 20/11/24  2:44 PM | 21/11/24  2:45 PM | 24 hr 1 min |
| 3 | 20/11/24  2:20 PM | 21/11/24  2:35 PM | 24 hr 15 min |
| 4 | 20/11/24  1:51 PM | 21/11/24  2:11 PM | 24 hr 20 min |
| 5 | 20/11/24  1:11 PM | 21/11/24  1:37 PM | 24 hr 26 min |
| 6 | 20/11/24  12:36 PM | 21/11/24  1:16 PM | 24 hr 40 min |

**Table S14. Otago Peninsula air sampling details for samples collected 18 days post-removal on 28/11/24.**

| **Site** | **Start** | **End** | **Duration** |
| --- | --- | --- | --- |
| 1 | 27/11/24  11:09 AM | 28/11/24  10:22 AM | 23 hr 13 min |
| 2 | 27/11/24  10:48 AM | 28/11/24  10:14 AM | 23 hr 26 min |
| 3 | 27/11/24  10:27 AM | 28/11/24  10:08 AM | 23 hr 41 min |

**Table S15. Otago Peninsula air sampling details for samples collected 25 days post-removal on 5/12/24.**

| **Site** | **Start** | **End** | **Duration** |
| --- | --- | --- | --- |
| 1 | 4/12/24  11:00 AM | 5/12/24  10:30 AM | 23 hr 30 min |
| 2 | 4/12/24  10:43 AM | 5/12/24  10:23 AM | 23 hr 40 min |
| 3 | 4/12/24  10:26 AM | 5/12/24  10:16 AM | 23 hr 50 min |

**Table S16. Otago Peninsula air sampling details for samples collected 32 days post-removal on 12/12/24.**

| **Site** | **Start** | **End** | **Duration** |
| --- | --- | --- | --- |
| 1 | 11/12/24  10:51 AM | 12/12/24  10:04 AM | 23 hr 13 min |
| 2 | 11/12/24  10:34 AM | 12/12/24  10:12 AM | 23 hr 38 min |
| 3 | 11/12/24  10:14 AM | 12/12/24  10:04 AM | 23 hr 50 min |

**Table S17. Otago Peninsula positive air sampling results including Cq value and DNA Quantity (copies/uL).**

| **Site** | **Technical replicates positive** | **Distance from source (metres)** | **Sample date** | **Cq** | **Quantity (copies/uL)** |
| --- | --- | --- | --- | --- | --- |
| 8 | 1/3 | 100 | 1 day post-deployment  31/10/24 | 37.37 | 21.42 |
| 1 | 3/3 | 1 | 6 days post-deployment  5/11/24 | 33.57 | 104.50 |
| 1 |  | 1 | 6 days post-deployment  5/11/24 | 34.05 | 77.05 |
| 1 |  | 1 | 6 days post-deployment  5/11/24 | 34.01 | 78.70 |
| 2 | 1/3 | 1 | 7 days post-deployment  6/11/24 | 35.54 | 29.68 |
| 5 | 1/3 | 10 | 8 days post-deployment  7/11/24 | 38.22 | 8.70 |
| 7 | 1/3 | 100 | 2 days post-removal  12/11/24 | 35.82 | 12.81 |
| 5 | 2/3 | 10 | 3 days post-removal  13/11/24 | 36.97 | 39.75 |
| 5 |  | 10 | 3 days post-removal  13/11/24 | 35.67 | 82.95 |
